# Epigenetic silencing of transposable elements by IRF2BP2 is a selective dependency of myeloid leukemia

**DOI:** 10.1101/2025.11.12.688028

**Authors:** Jingru Xu, Jessica Whittle, Liam Clayfield, Michael Lie-a-ling, Eve Augusta Leddy, Duncan Smith, Robert Sellers, Ozgen Deniz, Georges Lacaud

## Abstract

Acute myeloid leukemia (AML) is a heterogeneous malignancy with limited curative treatment options. Transposable elements (TEs) are now recognized as key regulators of genome function, with aberrant activation implicated in cancer. However, their tumor-type-specific roles remain poorly characterized. Using single-cell Perturb-seq, we systematically screened for chromatin-associated regulators in primary AML patient cells to uncover dependencies required for leukemia cell viability. Our screen identified IRF2BP2 as an AML-selective dependency, functioning as a repressor of TE expression. Loss of IRF2BP2 induced differentiation, apoptosis, and impaired leukemic cell fitness, phenotypes linked to transcriptional activation of TEs, particularly evolutionarily young human endogenous retrovirus K (HERV-K). Mechanistically, IRF2BP2 cooperates with TRIM28 and DNMT1 to epigenetically silence TE expression. CRISPR-mediated activation of HERV-K/LTR5_Hs recapitulated the phenotypic effects of IRF2BP2 loss, while targeted re-silencing of HERV-K/LTR5_Hs partially rescued the effects, establishing a causal link between TE regulation and AML maintenance. Our findings highlight tumor-suppressive functions of TEs in leukemia and reveal IRF2BP2 as a key regulator of TE silencing in AML. Targeting the epigenetic machinery governing TE repression may represent a promising therapeutic avenue for differentiation-inducing and immunomodulatory strategies in AML.

## Introduction

Acute myeloid leukemia (AML) is an aggressive hematologic malignancy characterized by a block of myeloid differentiation and uncontrolled expansion of myeloid progenitors. Conventional chemotherapy and allogeneic hematopoietic-stem-cell transplantation often have limited efficacy in treating AML^1^. The paucity of effective treatment options, coupled with adverse side effects and a high frequency of relapse, underscores the urgent need for novel therapeutic strategies to improve patient outcomes. AML is a malignancy whose subtypes are commonly classified based on distinct mutations and chromosomal translocations, a large proportion of which converge on epigenetic dysregulation^2^. These epigenetic aberrations play a critical role in enforcing differentiation blocks and sustaining leukemic cell survival, suggesting that targeting epigenetic dysregulation presents a promising therapeutic avenue in AML. However, the development of effective treatments remains a major challenge notably due to the heterogeneity of the disease. Identifying and characterizing shared dependencies on chromatin-associated proteins across AML subtypes, that are not directly altered by mutations or translocations, could reveal more broadly applicable therapeutic targets.

Approximately 50% of the human genome consists of transposable elements (TEs), which are commonly classified into two categories: Class I retrotransposons, which move via a copy-and-paste mechanism involving an RNA intermediate, and Class II DNA transposons, which move through a direct cut-and-paste mechanism^3,4^. Retrotransposons can be further separated into three major classes: (1) long terminal repeat (LTR) elements including endogenous retroviruses (ERVs); and non-LTR elements, which includes (2) long Interspersed Nuclear Elements (LINEs) and (3) short Interspersed Nuclear Elements (SINEs)^3,4^. Over evolutionary time, these TEs have contributed to genome expansion and variation^5,6^. Most of TEs are silenced in normal somatic cells and uncontrolled transposition of TEs can lead to genomic mutagenesis, instability, and dysregulation of gene expression^7,8^. Due to their viral-like characteristics, TEs can also elicit inappropriate inflammatory responses (viral mimicry), resulting in cellular stress, senescence, ageing or programmed cell death^9–15^. Therefore, TEs must be under strict epigenetic control to maintain genome integrity and avoid aberrant immune activation.

Tripartite Motif Containing 28 (TRIM28) and DNA Methyltransferase 1 (DNMT1) functionally cooperate in establishing and maintaining repressive chromatin states^16–18^ and are key regulators of TE silencing^13,19–26^. TRIM28 (KAP1 or TIF1β) is essential for silencing TEs during neural development and plays a critical role in repressing TE activity in embryonic stem cells^19–21,26^. DNMT1 is a DNA methyltransferase responsible for maintaining DNA methylation patterns upon replication. Recent studies have shown that DNMT1 also exhibits *de novo* methylation activity that specifically affects TEs^22^. Disruption of DNMT1, or treatment with epigenetic therapies, including hypomethylating agents targeting DNMTs, can induce TE activation, leading to innate antiviral immune responses including interferon signaling^9,10,13,22–24,27,28^. Pan-cancer genomic analyses have identified acquired retrotransposition events across a broad spectrum of cancer types. Notably, AML and myeloproliferative neoplasms exhibit the lowest incidences of retrotransposition^29^. In AML, LINE-1 retrotransposition has been proposed to exert a tumor-suppressive function^8^. Activation of TEs has also been shown to induce inflammatory responses that are detrimental to leukemic cell survival^15^. Consequently, epigenetic repression of retrotransposons is thought to confer a selective advantage for AML maintenance. Together, these findings suggest that leukemic cells require chromatin-associated proteins to repress cell intrinsic inflammatory responses driven by TE activation.

In recent years, the chromatin-associated protein Interferon Regulatory Factor 2 Binding Protein 2 (IRF2BP2) has emerged as a negative regulator of inflammatory responses in AML^30,31^. IRF2BP2 is a member of the IRF2BP protein family, which also includes IRF2BP1 and IRF2BPL. These family members share a conserved C-terminal RING domain and an N-terminal zinc finger domain, both of which are implicated in protein-protein interactions^32–35^. IRF2BP2 was initially identified as a transcriptional coregulator of Interferon Regulatory Factor 2 (IRF2)^32^. Beyond its interaction with IRF2, IRF2BP2 has been shown to exert IRF2-independent functions, regulating biological processes including cell survival, immunomodulation, differentiation, angiogenesis, and cell migration^33–40^. Here, we identify IRF2BP2 as a common dependency across a wide range of AMLs and reveal a previously uncharacterized mechanism by which IRF2BP2 controls cell-intrinsic inflammatory responses in AML. Our results show that IRF2BP2 knockout results in increased transcription of TEs. Mechanistically, this effect depends on IRF2BP2 associating with, and participating in maintaining, the TRIM28-DNMT1 complex. Loss of IRF2BP2 results in the dissociation of TRIM28-DNMT1 complex and subsequent activation of TE expression. These results position IRF2BP2 as a gatekeeper of TE activity in AML and highlight the IRF2BP2 axis as a promising therapeutic target across a broad range of AMLs.

## Results

### A Perturb-seq-based strategy reveals key regulators of AML

To identify key chromatin-associated factors in AML, we selected the top 15 candidates-excluding common essential genes-based on their CRISPR gene effect score across 24 AML cell lines in the DepMap database^41^ (Fig. 1a). To functionally validate these candidates in a more clinically relevant setting, we performed a CRISPR interference (CRISPRi)-based Perturb-seq screen^42^ in primary MLL-AF6 patient cells, a subtype of AML considered among the most aggressive AML and associated with poor prognosis^43^. Perturb-seq is a high-throughput single-cell CRISPR screening approach, enabling systematic analyses of the effects of specific genetic modifications at a single-cell level^44^. Compared to Cas9 nuclease-catalyzed DNA cleavage, CRISPRi minimizes the risk of genomic rearrangements and DNA damage-associated toxicity. This approach also increases the proportion of cells with uniform gene perturbation and enhances perturbation consistency, especially in primary samples^45-48^_._

**Fig. 1.**
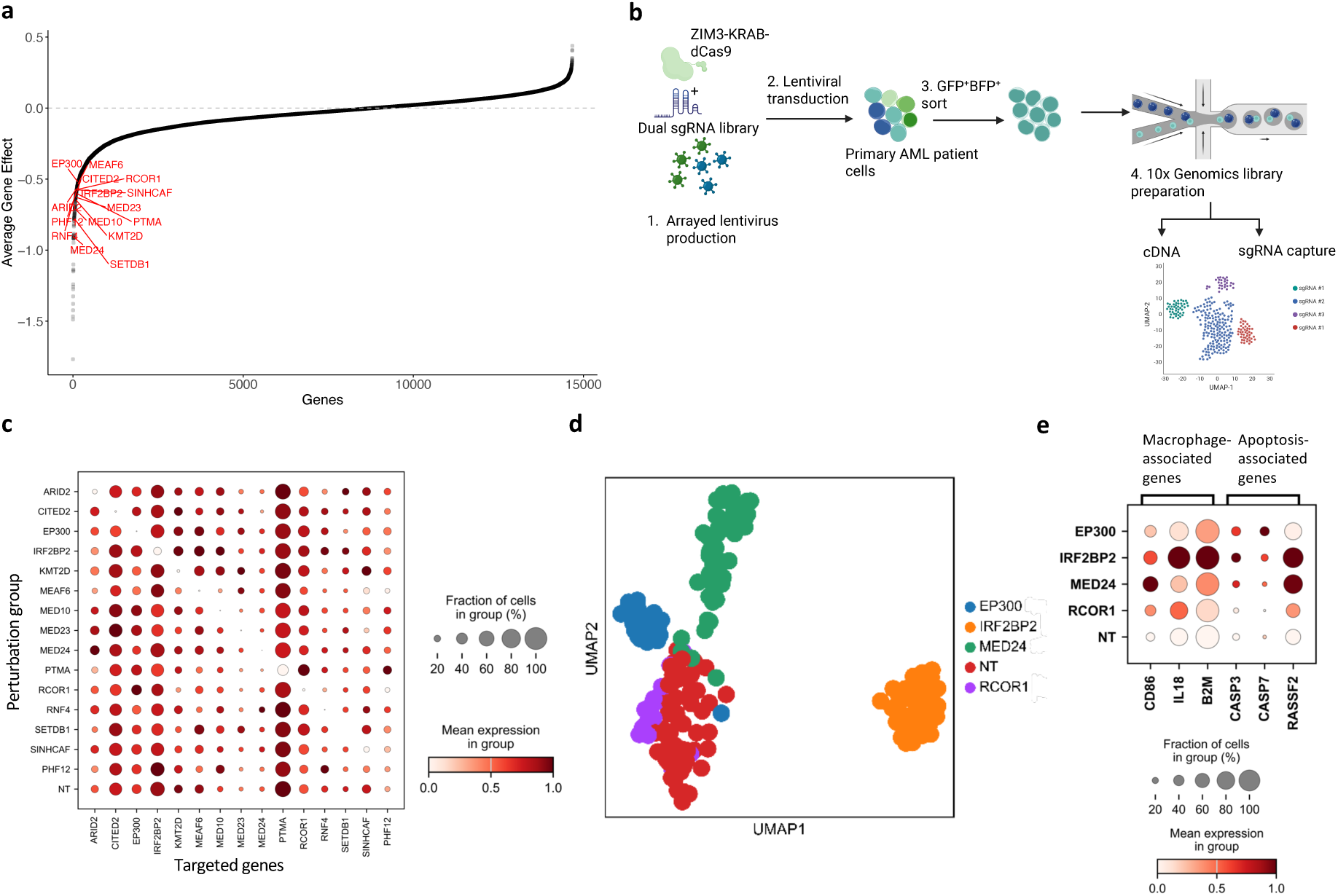
Single-cell perturb-seq identifies key chromatin-associated regulators in AML. **a**, Average CRISPR (chronos) gene effect scores across 24 AML cell lines in the DepMap database. Each point represents a single gene, ranked by its average Chronos gene effect score. The top 15 chromatin-associated regulators (highlighted in red), excluding common essential genes, were selected for perturb-seq. **b**, Schematic overview of the CRISPRi-based Perturb-seq experiment performed on primary human MLL-AF6 cells. **c**, Dotplot depicting the mean expression levels as well as the percentage of expressing cells for each targeted gene (x-axis) across all perturbation groups (y-axis), indicating the efficiency and specificity of knockdown (KD) for each targeted gene. **D**, UMAP projections generated using Mixscape for perturbation groups targeting *IRF2BP2, MED24, EP300, and RCOR1*, showing distinct perturbation signatures, especially for *MED24* and *IRFBP2*, compared to the NT control. **e**, Dot plot showing increased expression of macrophage-associated and apoptosis-associated genes (x-axis), along with the percentage of expressing cells, across perturbation groups (y-axis) compared to the NT control.

We constructed a lentiviral sgRNA library encoding a BFP fluorescent marker and dual sgRNAs targeting each of the 15 candidate genes, along with non-targeting (NT) control sgRNAs. This library was subsequently co-transduced with ZIM3-KRAB-dCas9-GFP into the primary MLL-AF6 patient cells. After six days, GFP⁺/BFP⁺ double-positive cells were sorted and subjected to 5′ single-cell RNA-sequencing (scRNA-seq) using the 10x Genomics platform (Fig. 1b). In total, we profiled 733 cells with an average of 3,496 transcripts per cell. With the exception of sgRNAs targeting *PHF12,* all targeted genes showed a significant downregulation (p < 0.05) by their corresponding sgRNAs compared to the NT control (Fig. 1c and Extended data Fig. 1a). We then applied Mixscape^49^ for cell classification, which improved accuracy by controlling for confounding variation in the primary sample. Perturbation signatures resulting from IRF2BP2, MED24, EP300, and RCOR1 knockdown (KD) were appreciably divergent from the NT control based on Mixscape analysis, and were all characterized by upregulation of genes associated with differentiation and apoptosis (Fig. 1d,e). This effect was most pronounced in the IRF2BP2 and MED24 groups, whose average gene expression profiles were the most distinct from the NT control (Fig. 1d, Extended Data Fig. 1b). Gene ontology (GO) analysis of cells perturbed with IRF2BP2 or MED24 revealed enrichment for pathways related to apoptosis and myeloid differentiation (Extended Data Fig. 1c,d).

Utilizing the bloodspot database^50^, we next assessed the expression level of IRF2BP2 and MED24 in AML patient samples relative to normal hematopoietic cells. *MED24* expression did not show appreciable differences between normal and malignant hematopoiesis (Extended data Fig. 2a). In contrast, *IRF2BP2* expression was significantly elevated across multiple AML subtypes compared to normal CD34^+^ hematopoietic stem and progenitor cells (Fig. 2a). Interestingly, several solid tumors also displayed upregulation of *IRF2BP2*, although to a lesser extent than that observed in AML (Extended data Fig. 2b)^51^. Consistent with these observations, DepMap data^41^ indicate a stronger requirement for IRF2BP2 in myeloid leukemia compared to solid cancer types (Extended data Fig. 2c). Taken together, these data suggest that IRF2BP2 is a key regulator of AML maintenance and prompted us to further characterize its role in AML.

**Fig. 2.**
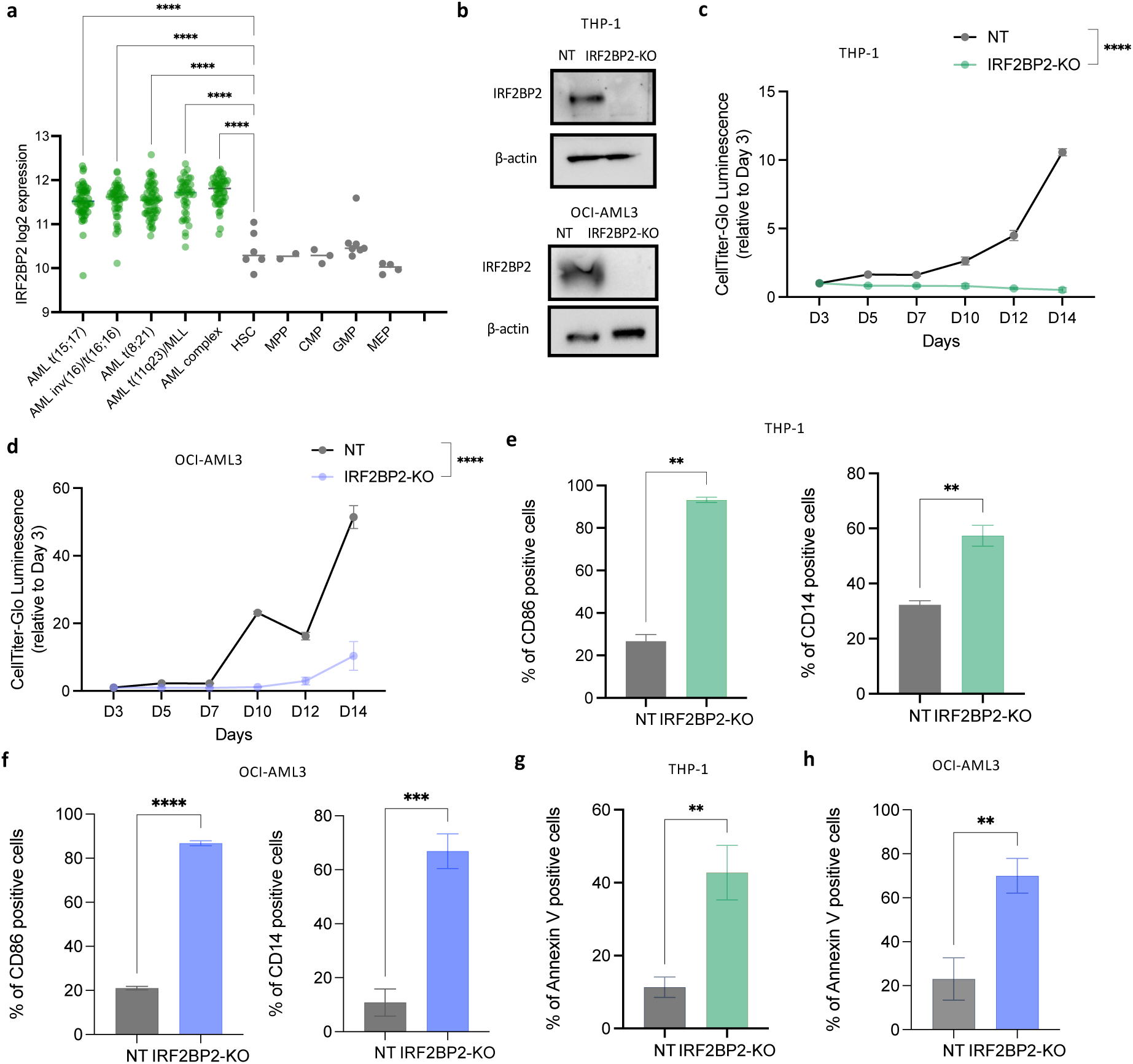
IRF2BP2 is a dependency across multiple AML subtypes. **a**, Gene expression levels of *IRF2BP2* were retrieved from the BloodSpot database. Mean *IRF2BP2* expression is significantly higher in AML patient samples (green dots) compared to normal hematopoietic stem and progenitor cells (grey dots). Each dot represents individual patient sample, horizontal lines indicate mean expression. Statistical test: one-way ANOVA. **b**, Western blot of IRF2BP2 following CRISPR/Cas9 sgRNA-mediated knockout (KO) in THP-1 (top) and OCI-AML3 (bottom) cell lines. β-actin was used as a loading control. **c,d**, Cell proliferation time course, as measured by CellTiter-Glo, for THP-1 (**c**) and OCI-AML3 (**d**) following CRISPR/Cas9-mediated KO of IRF2BP2. Transduced cells were sorted based on GFP and mCherry reporters on day 3. Statistical test: two-way ANOVA. **e,f**, Flow cytometry analysis of CD86 and CD14 expression 7 days post-transduction in THP-1 (**e**) and OCI-AML3 (**f**) cells. Statistical test: unpaired t-test. **g,h**, Flow cytometry analysis of apoptosis in THP-1 (**g**) and OCI-AML3 (**h**) 10 days post-transduction. Statistical test: unpaired t-test. **c-h**, Data shown as mean ± s.e.m (n = 3 biological experiments). ** *P* < 0.01; *** *P* < 0.001; **** *P* < 0.0001.

### IRF2BP2 is an essential vulnerability in both *in vitro* and *in vivo* AML models

To explore IRF2BP2 dependency in more detail, we next evaluated CRISPR-Cas9-based IRF2BP2 knockout (KO) in a panel of human AML cell lines including OCI-AML3 (NPM1^mut^), THP-1 (MLL-AF9), ML-2 (MLL-AF6), P31/Fujioka (MLLT10-PICALM), and OCI-AML2 (MLL-AF6). Two sgRNAs targeting *IRF2BP2* were used simultaneously to disrupt the locus and their effectiveness was confirmed by western blot in both THP-1 and OCI-AML3 cells (Fig. 2b). In all lines, IRF2BP2 KO led to a marked reduction in cell growth, along with significant induction of both differentiation (indicated by increased CD86 and CD14 expression) and apoptosis, independently of the genetic aberration in the cell lines (Fig. 2c–h and Extended data Fig. 3a–i). To confirm that the observed phenotypic effects were specifically attributable to IRF2BP2 depletion, we reintroduced IRF2BP2 into IRF2BP2-KO cells. This rescue reversed the alterations in cell differentiation and apoptosis (Extended Data Fig. 4a–d). Together, these findings demonstrate that IRF2BP2 is essential for the survival of AML cells across diverse genetic backgrounds, suggesting a broad role for IRF2BP2 in supporting leukemic cell viability.

To further investigate the role of IRF2BP2 in leukemogenesis, we extended our analysis to *in vivo* models. Human AML cell lines were co-transduced with Cas9 and either IRF2BP2-targeting sgRNAs or NT sgRNAs, and subsequently transplanted into immunodeficient recipient mice. IRF2BP2 KO in OCI-AML3 cells resulted in markedly prolonged survival and reduced leukemia burden, with all recipients surviving beyond 100 days, in contrast to the rapid disease onset observed in control mice (Fig. 3a,b). Similarly, THP-1 cells depleted of IRF2BP2 showed significantly delayed leukemia progression and reduced leukemia burden, with one long-term survivor exceeding 100 days post-transplantation (Fig. 3c,d). Notably, in mice that developed leukemia, sorted CD45⁺ human AML cells transduced with IRF2BP2-targeting sgRNAs retained IRF2BP2 protein expression comparable to control cells (Fig. 3e), indicating the outgrowth of escaper cells and suggesting strong selective pressure against IRF2BP2 loss *in vivo*. Moreover, IRF2BP2 depletion in additional AML cell lines (ML-2, OCI-AML2, and P31/Fujioka) similarly resulted in significantly delayed leukemia progression (Fig. 3f–h). Collectively, these data demonstrate that IRF2BP2 is essential for AML progression and that its loss impairs leukemogenesis *in vivo*.

**Fig. 3.**
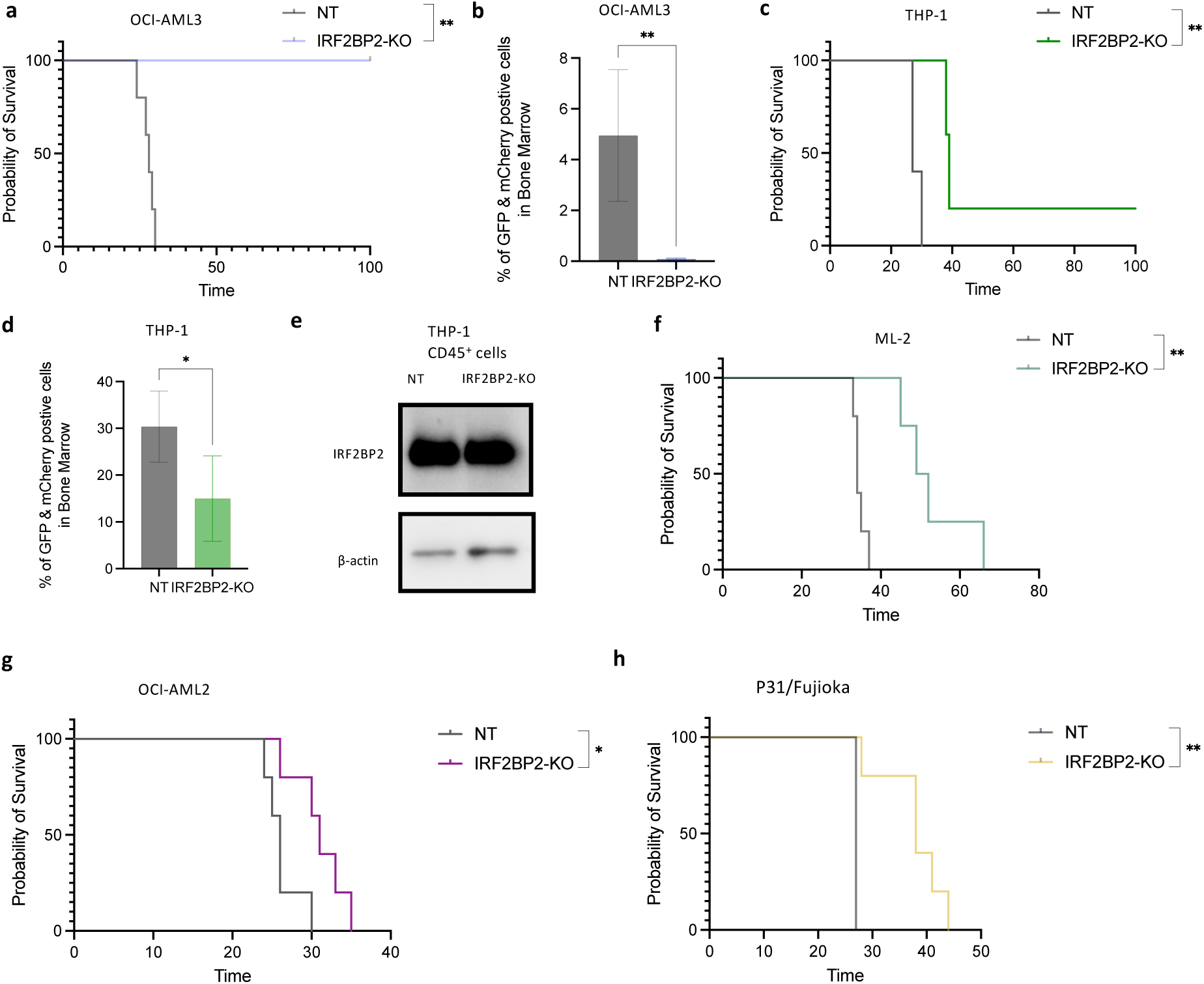
IRF2BP2 depletion significantly prolongs survival and reduces leukemia burden *in vivo*. **a**, Kaplan–Meier survival curves of NSGS mice xenografted with OCI-AML3 cells transduced with either NT sgRNAs or IRF2BP2-targeting sgRNAs. N = 5 mice per group, Log-rank Mantel-Cox test. **b**, Flow cytometry analysis of leukemic burden in bone marrow aspirates from sublethally irradiated NSGS mice transplanted with transduced OCI-AML3 cells. Results are the mean ± s.e.m. (n = 5) and analyzed by unpaired t-test**. c**, Kaplan–Meier survival curves of NSGS mice transplanted with THP-1 cells transduced with either NT sgRNAs or IRF2BP2-targeting sgRNAs. N = 5 mice per group, Log-rank Mantel-Cox test. **d**, Flow cytometry analysis of leukemic burden in bone marrow aspirates from mice transplanted with transduced THP-1 cells. Results are the mean ± s.e.m. (n = 5) and analyzed by unpaired t-test**. e**, Western blot of IRF2BP2 in human CD45^+^ cells from mice transplanted with THP-1 cells that developed AML, indicating the outgrowth of cells that escaped Cas9-mediated KO of IRF2BP2. β-actin was used as a loading control. **f–h**, Kaplan–Meier survival curves of NSGS mice transplanted with ML-2 (**f**), OCI-AML2 (**g**) or P31/Fujioka (**h**) cells transduced with either NT sgRNAs or IRF2BP2-targeting sgRNAs. N = 5 mice per group, Log-rank Mantel-Cox test. * *P* < 0.05; ** *P* < 0.01.

### IRF2BP2 interacts with TRIM28 and DNNMT1 in AML

IRF2BP2 is a transcriptional co-factor that likely exerts its function through interactions with other chromatin-associated proteins. To elucidate the molecular mechanisms underlying its leukemia-specific role, we sought to identify endogenous IRF2BP2-interacting partners in AML using Rapid Immunoprecipitation Mass Spectrometry of Endogenous Proteins (RIME). To identify high-confidence conserved interactors, we performed IRF2BP2-RIME in five distinct human AML cell lines (THP-1, OCI-AML2, ML-2, P31/Fujioka, and OCI-AML3), each in triplicate, using stringent criteria (Fig. 4a). For each cell line, we retained only those proteins consistently detected across the three independent IRF2BP2-RIME replicates and excluded any proteins identified in matched IgG controls. We then intersected the resulting interactomes across all five AML lines, revealing a conserved core set of IRF2BP2-associated proteins, including IRF2BP2 itself, IRF2BP1, TRIM28, DNMT1, NCL, EIF4A3, and RPL7 (Fig. 4b, and Supplementary Table 2). The identification of IRF2BP1, a known IRF2BP2 binding partner^30,34^, confirms the specificity and robustness of our approach. Several of the newly identified interactors are key regulators of chromatin dynamics. Notably, TRIM28, a multifunctional transcriptional regulator involved in chromatin remodeling, and DNMT1, an essential DNA that maintains DNA methylation patterns, are known to function as a complex mediating epigenetic silencing, particularly of TEs^13,19–25^ (Fig. 4c)^52^. Our data identify IRF2BP2 as a previously unrecognized interactor of the TRIM28–DNMT1 complex, suggesting a cooperative role for IRF2BP2 in chromatin-associated gene regulation in AML.

**Fig. 4.**
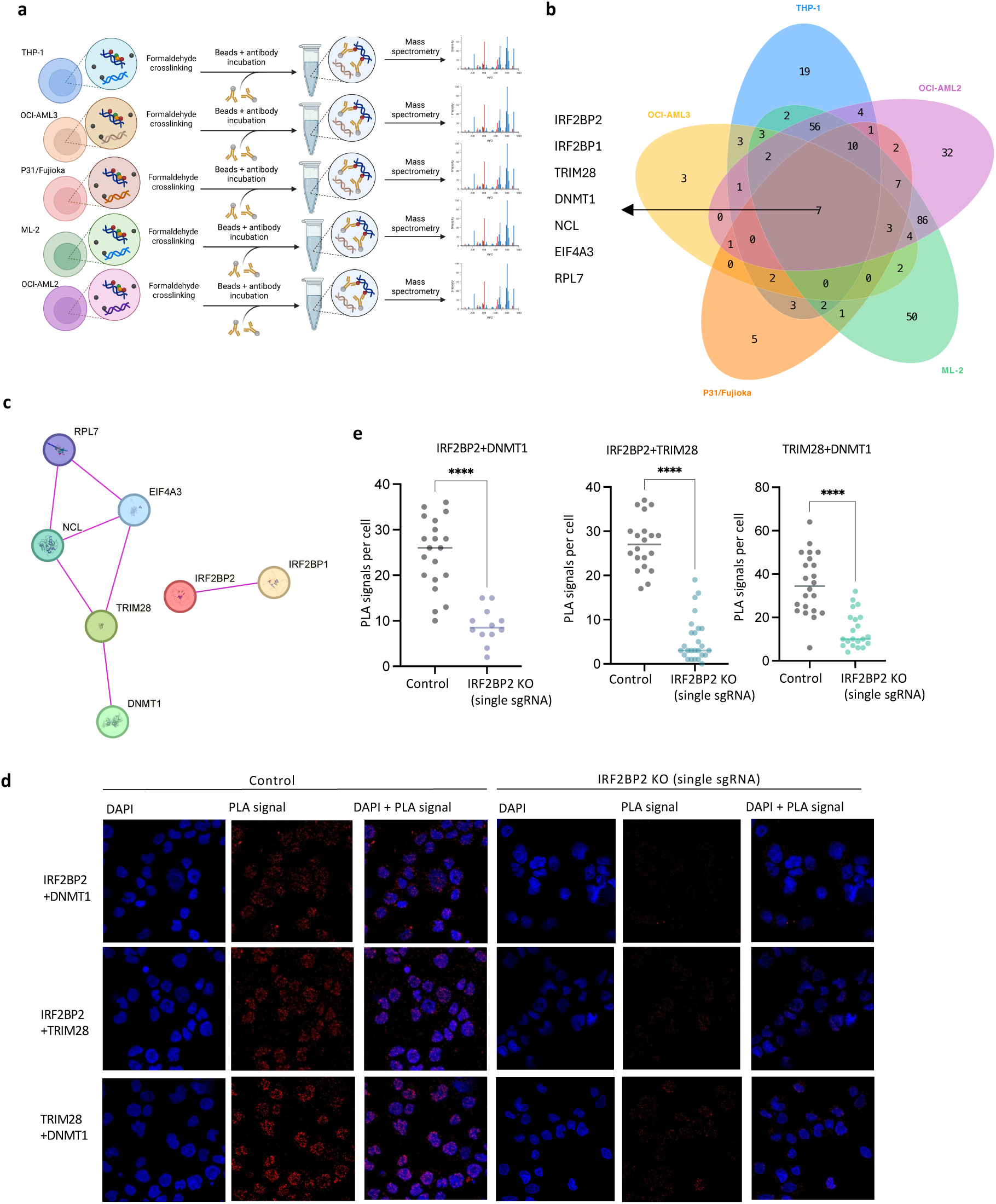
IRF2BP2 interacts with TRIM28 and DNMT1 in AML. **a**, Schematic of Rapid Immunoprecipitation Mass Spectrometry of Endogenous Proteins (RIME) of IRF2BP2 in 5 human AML cell lines. **b**, IRF2BP2-RIME was performed in THP-1, OCI-AML3, P31/Fujioka, OCI-AML2, and ML-2 cells with three biological replicates per cell line. The Venn diagram shows a conserved core set of IRF2BP2-associated proteins across the 5 cell lines. **c**, Protein–protein interaction network of the intersecting IRF2BP2-associated proteins identified by RIME, generated using the STRING database. Nodes represent proteins, and lines represent known interactions. The network shows that IRF2BP2 has not previously identified as an interactor of TRIM28 or DNMT1 based on the STRING database. **d,e**, Duolink Proximity Ligation Assay (PLA) was used to validate the interaction between IRF2BP2, TRIM28 and DNMT1 in THP-1 cells. Negative control cells or IRF2BP2 single-sgRNA KO cells were stained with the corresponding antibodies, and fluorescent signals was measured using Zeiss confocal microscope. Fluorescent PLA signals (red dots) indicate close spatial proximity between the two proteins, consistent with a protein–protein interaction. Loss of IRF2BP2 reduced the detectable interaction between TRIM28 and DNMT1 (bottom panels of **d**; right panel of **e**). Nuclei were detected by DAPI (blue). Unpaired t-test. **** *P* < 0.0001.

We next validated the interactions between IRF2BP2, TRIM28, and DNMT1 using Duolink proximity ligation assays (PLA). In THP-1 cells, we detected specific interactions between IRF2BP2 and TRIM28, IRF2BP2 and DNMT1, as well as TRIM28 and DNMT1 (Fig. 4d,e). As expected, IRF2BP2 KO with single sgRNA targeting *IRF2BP2* resulted in a significant reduction in PLA signals between IRF2BP2 and TRIM28, as well as between IRF2BP2 and DNMT1 (Fig. 4d,e). Notably, IRF2BP2 KO also led to a marked decrease in the interaction signals between TRIM28 and DNMT1, suggesting that IRF2BP2 not only binds to both proteins but also contributes to stabilizing their association. Together, these data demonstrate that IRF2BP2 is a component of the TRIM28–DNMT1 epigenetic regulatory complex and plays a critical role in maintaining its stability.

### Perturbation of IRF2BP2 interactors TRIM28 and DNMT1 recapitulates the phenotypes observed upon IRF2BP2 KO

Having identified IRF2BP2 as a novel component of the TRIM28–DNMT1 complex, we next assessed whether DNMT1 and TRIM28 play a functional role similar to that of IRF2BP2. To test this, we first inhibited DNMT1 using GSK3685032, a selective DNMT1 inhibitor that reduces both DNMT1 protein levels and enzymatic activity^27^. DNMT1 protein levels decreased in a dose-dependent manner following treatment (Extended Data Fig. 5a). GSK3685032 treatment led to a significant reduction in cell proliferation, along with markedly increased differentiation and apoptosis in both THP-1 and OCI-AML3 cell lines (Fig. 5a–f), closely recapitulating the phenotypes observed with IRF2BP2 KO. To further explore the relevance of this complex, we depleted TRIM28 using a CRISPR-Cas9 strategy with two independent sgRNAs and confirmed KO efficiency by Western blot (Extended Data Fig. 5b).

**Fig. 5.**
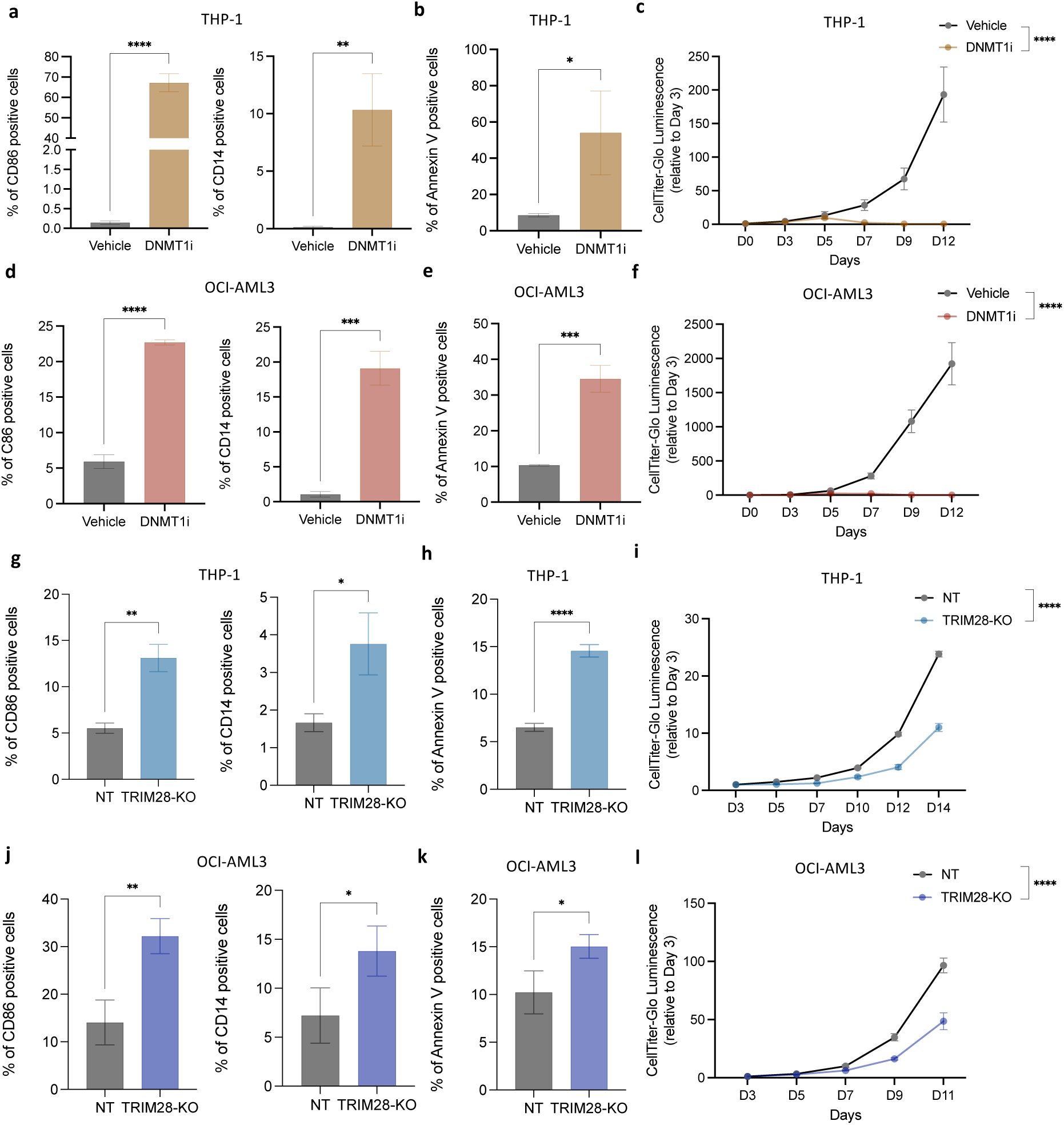
Depletion of DNMT1 or TRIM28 replicates the phenotype observed in IRF2BP2 KO cells. a-f. Phenotypic analyses of THP-1 and OCI-AML3 cells treated with 400nM GSK3685032 (DNMT1 inhibitor). **a,d**, Flow cytometry analysis of myeloid differentiation markers CD14 and CD86 in THP-1 (**a**) and OCI-AML3 (**d**) cells after 7 days treatment with GSK3685032. Unpaired t-test. **b,e**, Flow cytometry analysis of apoptosis induction following GSK3685032 treatment relative to control in THP-1 (**b**) and OCI-AML3 (**e**) cells after 7 days of treatment. Unpaired t-test. **c,f**, CellTiter-Glo assay assessing the effect of GSK3685032 treatment on the proliferation of THP-1 (**c**) and of OCI-AML3 (**f**) cells. Two-way ANOVA. **g-l**, phenotypic analysis of THP-1 and OCI-AML3 cells following CRISPR/Cas9-mediated KO of TRIM28. **g,j**, Cell differentiation was assessed in THP-1 (**g**) and OCI-AML3 (**j**) cells 7 days post-transduction by flow cytometry. Unpaired t-test. **h,k**, Flow cytometry analysis of apoptosis in THP-1 (**h**) and OCI-AML3 (**k**) cells 10 days post-transduction. Unpaired t-test. **i,l**, Cell proliferation was assessed in THP-1 (**i**) and OCI-AML3 (**l**) cells 10 days post-transduction using CellTiter-Glo assay. Two-way ANOVA. Data shown as mean ± s.e.m (n = 3). * *P* < 0.05; ** *P* < 0.01; *** *P* < 0.001; **** *P* < 0.0001.

TRIM28 loss resulted in a more modest, yet significant, increase in cell differentiation and apoptosis, as well as decreased proliferation (Fig. 5g–l). Together, these findings suggest that depletion of DNMT1 or TRIM28 phenocopies the effects of IRF2BP2 loss, suggesting a coordinated role for all three proteins in supporting AML cell survival.

### IRF2BP2 is required to suppress the anti-viral immune response triggered by TE activation

To uncover the underlying mechanism by which IRF2BP2 complex regulates AML survival, we performed RNA sequencing on THP-1 cells and OCI-AML3 cells following IRF2BP2 KO (Fig. 6a,b). Loss of IRF2BP2 in both cell lines led to upregulation of immune-related pathways, including positive regulation of cytokine production and inflammatory responses (Extended Data Fig. 6a,b), consistent with prior studies^30,31^. Over 65% of genes upregulated and 53% of genes downregulated upon IRF2BP2 loss in OCI-AML3 cells overlapped with those altered in THP-1 cells (Fig. 6c,d). Notably, the top shared upregulated signatures included pathways related to the defense response to viruses (Fig. 6e). This antiviral response may be linked to the activation of TEs, as both DNMT1 and TRIM28—known binding partners of IRF2BP2—have previously been implicated in TE silencing.

**Fig. 6.**
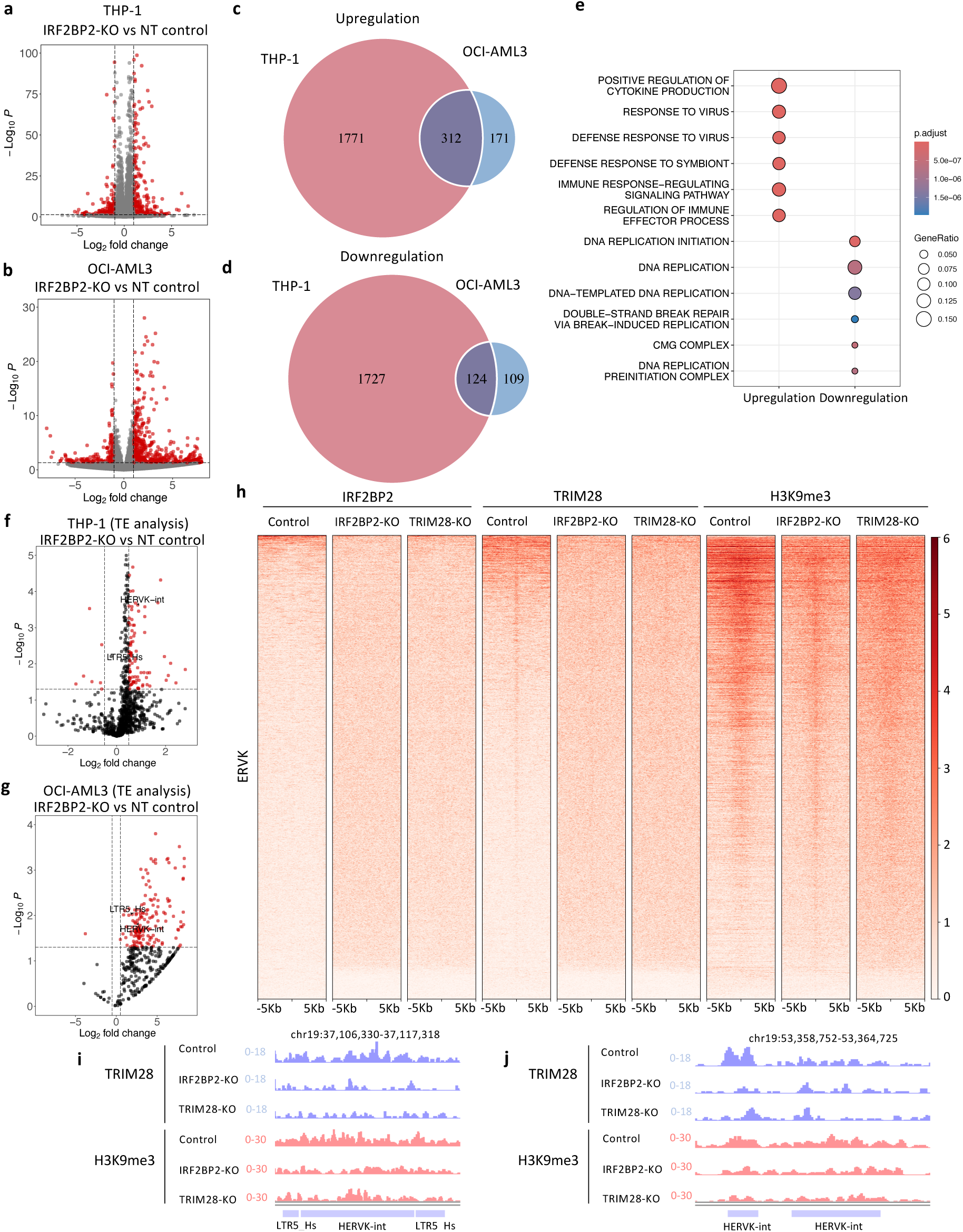
Loss of IRF2BP2 induces the antiviral immune response. a-g. RNA-seq of IRF2BP2 KO and NT control in THP-1 and OCI-AML3 cells. Volcano plot showing differentially expressed genes in THP-1 (**a**) and OCI-AML3 (**b**) cells at 3 days post-transduction. Significantly upregulated and downregulated genes (P < 0.05, |log₂ fold change| > 1) are highlighted in red. **c,d**, Venn diagram illustrating the overlap of significantly upregulated genes (**c**) and downregulated genes (**d**) (adjusted P < 0.05) between THP-1 cells and OCI-AML3 cells upon IRF2BP2 depletion. **e**, Gene Ontology (GO) analysis illustrating enriched biological processes among intersecting differentially expressed genes following IRF2BP2 KO in THP-1 and OCI-AML3 cells. **f,g**, Transposable element analysis in IRF2BP2-KO versus NT control cells in THP-1 (**f**) and OCI-AML3 (**g**). Significantly upregulated and downregulated elements (P < 0.05, |log₂ fold change| > 0.5) are highlighted in red. **h**, Heatmap plots of CUT&RUN data for IRF2BP2, TRIM28 and H3K9me3 at ERVK regions across control, IRF2BP2-KO and TRIM28-KO THP-1 cells. **i,j**, Genome browser image illustrating reduced TRIM28 and H3K9me3 occupancy at HERV-K/LTR5_Hs elements following loss of IRF2BP2 and TRIM28.

To assess whether loss of IRF2BP2 triggers viral defense response via TE upregulation, we analyzed TE expression following IRF2BP2 KO. Perturbation of IRF2BP2 led to significant upregulation of TEs compared to NT controls in both THP-1 and OCI-AML3 cells: IRF2BP2 depletion upregulated 249 TE sub-families in THP-1, and 159 in OCI-AML3 (Fig. 6f,g). Among those upregulated TEs, members of the ERVK family, HERV-K (HML-2) and its regulatory promoter LTR5_Hs, were co-upregulated in both cell lines. HERV-K is the youngest and most active family of human endogenous retrovirus and is capable of triggering antiviral immune responses^53,54^. Moreover, analysis of a published RNA-seq dataset from THP-1 cells treated with the DNMT1 inhibitor GSK3685032^27^ revealed similar transcriptional changes, including increased expression of cytokine and inflammatory genes upon DNMT1 loss, consistent with the effects observed following IRF2BP2 depletion (Extended Data Fig. 7a,b). Over 40% of genes dysregulated upon IRF2BP2 KO overlapped with those altered by DNMT1 inhibition and the top shared gene signature was enriched for the defense response to virus pathway (Extended Data Fig. 7c). Consistent with the effects of IRF2BP2 depletion, loss of DNMT1 resulted in broad activation of TEs (Extended Data Fig. 7d). These findings further support a coordinated role of the IRF2BP2 complex in suppressing TE expression and limiting antiviral immune responses.

To validate IRF2BP2 is a key regulator of ERVK expression, we next performed CUT&RUN for IRF2BP2, TRIM28, and H3K9me3 in control, IRF2BP2-KO and TRIM28-KO THP-1 cells. Depletion of IRF2BP2 or TRIM28 markedly disrupted the binding of both TRIM28 and H3K9me3 at TE regions (Fig. 6h). Although IRF2BP2 itself does not consistently bind all TE regions occupied by TRIM28 and H3K9me3, our occupancy analysis consistently showed a marked reduction in TRIM28 and H3K9me3 occupancy at TE loci, including HERV-K/LTR5_Hs, following IRF2BP2 depletion (Fig. 6h–j). In addition, loss of IRF2BP2 or TRIM28 also reduced TRIM28 and H3K9me3 occupancy at non-repetitive genomic regions (Extended Data Fig. 8), further supporting a critical role for IRF2BP2 in preserving heterochromatin structure and facilitating epigenetic regulation through the TRIM28–DNMT1 complex. Together with our Duolink PLA assays, which demonstrated that IRF2BP2 KO disrupted the physical interaction between TRIM28 and DNMT1 (Fig. 4d,e), these findings suggest that IRF2BP2 facilitates the assembly and/or stabilization of the TRIM28-DNMT1 complex and hence plays a critical role in repressing TE activity.

### Activation of HERV-K/LTR5_Hs impairs AML cell surival

To investigate the effect of ERVK activation on leukemogenesis, we used CRISPR activation (CRISPRa) to induce HERV-K/LTR5_Hs expression in THP-1 and OCI-AML3 cells. CARGO arrays containing sgRNAs targeting LTR5_Hs, with scaffolds derived from either *Streptococcus pyogenes* (LTR5_Hs-Sp) or *Staphylococcus aureus* (LTR5_Hs-Sa)^55^, were employed to drive expression of both LTR5_Hs and HERV-K. Cells were first engineered to stably express either LTR5_Hs-Sp or LTR5_Hs-Sa sgRNAs via the PiggyBac transposon system, followed by delivery of the dCas9-VP64 activator (Extended Data Fig. 9a). As dCas9-VP64 binds exclusively to the *S. pyogenes* scaffold and not to the *S. aureus* scaffold, the LTR5_Hs-Sa group served as a negative control. Elevated expression of HERV-K/LTR5_Hs in the LTR5_Hs-Sp group, compared to the LTR5_Hs-Sa negative control, was confirmed by RT-qPCR in both THP-1 and OCI-AML3 cells (Extended Data Fig. 9b,c). We observed increased cellular differentiation, enhanced apoptosis, and impaired proliferation following CRISPR-mediated activation of HERV-K/LTR5_Hs in both THP-1 and OCI-AML3 cells (Fig. 7a–f), consistent with the phenotypic effects observed upon IRF2BP2 loss. In addition, activation of HERV-K/LTR5_Hs led to upregulation of genes involved in inflammatory responses (OAS1, OAS2, IFIT1, IFIT2 and GBP2), suggesting that TE activation triggers antiviral immune signaling pathways (Fig. 7g–h). Collectively, these findings support a functional connection between TE activation and reduced AML cell viability, underscoring the potential therapeutic relevance of targeting epigenetic regulators that silence TEs.

**Fig. 7.**
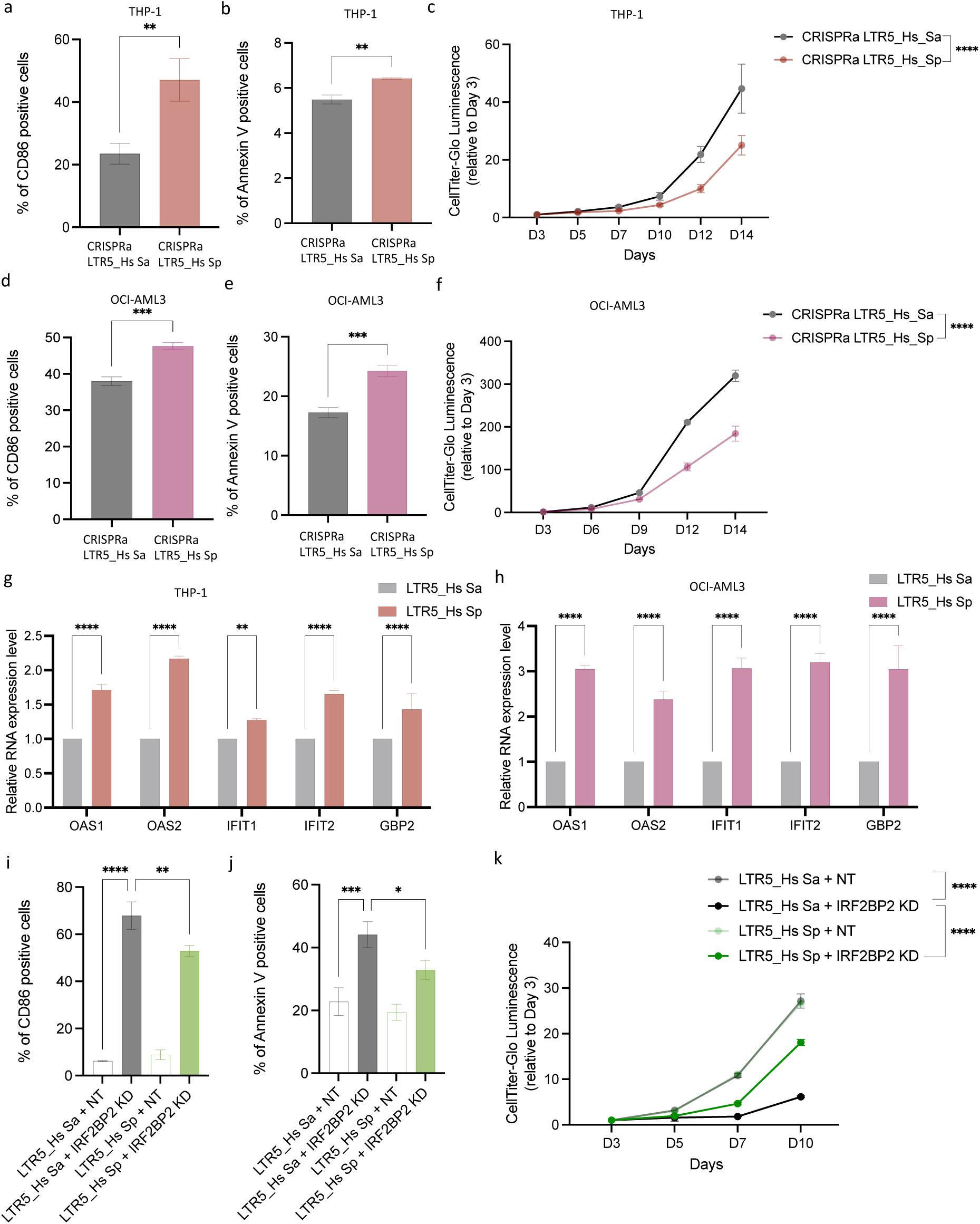
Activation of HERV-K/LTR5_Hs impairs AML cell survival, and repression of HERV-K/LTR5_Hs partially restores the reduced viability caused by IRF2BP2 depletion. a-h. Upregulation of HERV-K/LTR5_Hs in THP-1 and OCI-AML3 cells using CRISPR-mediated activation. **a,d**, Expression of the differentiation marker CD86 was assessed by flow cytometry in THP-1 (**a**) and OCI-AML3 (**d**) cells 7 days post-transduction following activation of HERV-K/LTR5_Hs. Unpaired t-test. **b,e**, Apoptotic cell populations were evaluated in THP-1 (**b**) and OCI-AML3 (**e**) cells by flow cytometry 10 days post-transduction using Annexin V and Hoechst staining. Unpaired t-test. **c,f**, Cell proliferation in THP-1 (**c**) and OCI-AML3 (**f**) cells was monitored over time using the CellTiter-Glo assay following HERV-K/LTR5_Hs activation. Two-way ANOVA. **g,h**, Activation of HERV-K/LTR5_Hs led to increased expression of genes associated with antiviral immune responses in THP-1 (**g**) and OCI-AML3 (**h**) cells, as determined by RT-qPCR analysis 4 days post-transduction. **i-k**, Rescue experiments using CRISPR-mediated interference were performed in OCI-AML3 cells. Cells were first transfected with either the LTR5_Hs *Staphylococcus aureus* scaffold CARGO array (LTR5_Hs Sa) or the LTR5_Hs *Streptococcus pyogenes* scaffold CARGO array (LTR5_Hs Sp), followed by transduction with dCas9-ZIM3-KRAB and sgRNAs targeting either *IRF2BP2* (IRF2BP2 KD) or a NT control. **i**, Cell differentiation was assessed by CD86 expression 7 days after transduction. One-way ANOVA. **j**, Apoptosis was evaluated 10 days after transduction using Annexin V staining. One-way ANOVA. **k**, Cell proliferation was measured using the CellTiter-Glo luminescence assay. Two-way ANOVA. Data shown as mean ± s.e.m (n = 3). * *P* < 0.05; ** *P* < 0.01; *** *P* < 0.001; **** *P* < 0.0001.

### Repression of HERV-K/LTR5_Hs partially rescues the IRF2BP2 KO phenotype in AML

We next tested whether the phenotypic effects of IRF2BP2 depletion could be rescued by repressing HERV-K/LTR5_Hs expression in OCI-AML3 cells. OCI-AML3 cells were selected because they exhibited higher HERV-K/LTR5_Hs expression upon IRF2BP2 loss compared to THP-1 cells (Fig. 6f.g). To this end, we used a CRISPR interference (CRISPRi) system based on *Streptococcus pyogenes* ZIM3-KRAB-dCas9 in combination with sgRNAs targeting *IRF2BP2* or a NT control. To specifically repress HERV-K/LTR5_Hs, OCI-AML3 cells were first engineered to stably express LTR5HS-Sa or LTR5HS-Sp sgRNAs using the PiggyBac transposon system. These cells were then transduced with lentiviral vectors encoding ZIM3-KRAB-dCas9 and either IRF2BP2-targeting or NT control sgRNAs (Extended Data Fig. 9d). Expression of LTR5HS-Sa or LTR5HS-Sp alone in the NT control background had no significant impact on cell phenotype, indicating that expression of these sgRNAs by themselves does not affect cell viability. As expected, IRF2BP2 depletion induced differentiation, increased apoptosis, and impaired proliferation. Notably, these effects were significantly attenuated in the LTR5HS-Sp group, where repression of HERV-K/LTR5_Hs partially rescued the IRF2BP2-depletion phenotype by reducing differentiation and apoptosis and restoring proliferative capacity (Fig. 7i–k).These findings suggest that IRF2BP2 maintains AML cell fitness, at least in part, by repressing HERV-K/LTR5_Hs expression, supporting the finding that TE activation is the key downstream consequence of IRF2BP2 loss that contributes to leukemia differentiation and apoptosis.

## Discussion

Chromatin-associated factors have emerged as promising therapeutic targets in AML due to their central role in regulating gene expression programs through genetic and epigenetic mechanisms that drive leukemogenesis. To effectively target AML across subtypes, a deeper understanding of the common epigenetic pathways that are dysregulated in the disease is essential. Recent studies have highlighted the importance of tight regulation of cell-intrinsic inflammatory responses in AML^30,31^. Notably, the activation of TEs—which comprise nearly half of the human genome—has emerged as a potent intrinsic trigger of inflammation^9–15^. Therefore, elucidating the epigenetic mechanisms by which AML cells suppress TE activation is of significant interest and may uncover therapeutic vulnerabilities.

Here, we identify the chromatin-associated factor IRF2BP2 as a dependency across multiple AML subtypes. We demonstrate that IRF2BP2 depletion in AML induces a cell-intrinsic inflammatory cascade, resulting in apoptosis, impaired proliferation, and reduced leukemia burden *in vivo*. These findings are consistent with previous studies that identified IRF2BP2 as an AML dependency involved in transcriptional repression of inflammatory response genes^30,31^. Our current study uncovers a novel critical role for IRF2BP2 in the genome-wide repression of TEs, which are potent activators of cell-intrinsic inflammatory responses. Mechanistically, we found that IRF2BP2 is an essential co-factor required to maintain a TRIM28- and DNMT1-containing repressive complex. Prior studies have established that both TRIM28 and DNMT1 are critical for TE silencing^13,19–26^. Our data support a model in which loss of IRF2BP2 disrupts the TRIM28–DNMT1 complex, resulting in widespread TE activation. This, in turn, triggers a robust inflammatory and viral mimicry response that compromises AML cell fitness. Notably, the elevated expression of IRF2BP2 in AML compared to normal hematopoietic cells suggests that AML cells co-opt this factor to enforce epigenetic suppression of TEs, thereby buffering the oncogenic stress imposed by mutant or rearranged epigenetic regulators.

IRF2BP2 expression is markedly elevated in AML compared to normal hematopoietic cells, representing the most pronounced tumor-specific upregulation among the cancer types analyzed. However, multiple solid cancers, including those of the breast, colon, and liver, also show increased IRF2BP2 expression relative to their matched normal tissues (Extended data Fig. 2b). This raises the possibility that IRF2BP2 may play broader roles in tumorigenesis, possibly employing the same epigenetic mechanism we characterized in AML. Interestingly, analysis of publicly available immunotherapy datasets via the ROC Plotter platform reveals that immunotherapy-resistant patient samples exhibit higher IRF2BP2 expression in a pan-cancer context^56^. This observation suggests that IRF2BP2 could potentially serve as a prognostic biomarker for immunotherapy-resistance across multiple cancer types. Although these datasets are derived from solid tumors, the consistent upregulation of IRF2BP2 in resistant cases raises the possibility that similar mechanisms may be relevant in AML. Activation of TEs represents one of the major sources of neoantigen production across multiple cancer types^57^. TE-derived dsRNAs, peptides, or chimeric proteins have the potential to stimulate T cell–mediated immune responses and may be leveraged in combination with therapeutic cancer vaccines or CAR-T cell therapies^58^. Perturbation of Trim28 in mouse models has previously been shown to sensitize cancers to immunotherapy, including PD-1 blockade^59^. In addition, treatment with DNMT inhibitor has been reported to induce neopeptides in AML patient samples^28^. Collectively, these findings imply that activation of TEs, achieved by targeting IRF2BP2 and its associated protein complexes, may represent a promising therapeutic avenue for enhancing neoantigen presentation and improving responses to immunotherapy.

Beyond their involvement in cancer, TEs have also been linked to ageing. TEs are reactivated with age due to the loss of epigenetic silencing mechanisms, which in turn triggers innate immune responses, promoting inflammation and age-related pathologies^14,60^. Analysis of single-cell transcriptomic data from the Tabula Muris Senis project revealed an age-associated decline in Irf2bp2 expression in murine white blood cells^61^, suggesting that Irf2bp2 may play a role in immune aging. This observed reduction in IRF2BP2 expression alongside age-associated TE upregulation provides insight into a potentially broader role for IRF2BP2 that extends beyond cancer to other contexts, such as ageing.

In this study, we employed RIME, which allows for the identification of protein-protein interactions on chromatin under native expression conditions^62^, to identify IRF2BP2-associated chromatin complex. It involves a formaldehyde crosslinking, which enhances the capture of low-abundance proteins and transient interactions that are often missed by conventional immunoprecipitation mass spectrometry approaches. In addition to TRIM28 and DNMT1, our RIME analysis identified several candidate RNA-binding proteins as novel IRF2BP2 interactors, including nucleolin (NCL), ribosomal protein L7 (RPL7), and eukaryotic initiation factor 4A-III (EIF4A3). NCL is a nucleolar phosphoprotein involved in rRNA transcription and ribosome assembly; RPL7 serves as a structural component of the ribosome; and EIF4A3 contributes to ribosome biogenesis through its role in mRNA processing^63–65^. Together, these data suggest a potential previously unappreciated role for IRF2BP2 in coordinating ribosomal function and biogenesis in leukemic cells.

In summary, our study identifies IRF2BP2 as a critical dependency in AML. Mechanistic investigations reveal that IRF2BP2, in cooperation with TRIM28 and DNMT1, suppresses TE expression and restrains aberrant immune activation, thereby preserving AML cell fitness (Extended Data Fig. 10). These findings provide novel insights into the epigenetic and immunoregulatory functions of IRF2BP2 in leukemia and lay a strong rationale for the development of therapeutic strategies targeting IRF2BP2 and its associated protein complexes.

## Methods

### Cells and cell culture

Human leukemia cell lines (THP-1, ML-2, and P31/Fujioka) were cultured in 90% RPMI (Gibco) with 10% FBS and 1% penicillin-streptomycin (Sigma-Aldrich). OCI-AML2 was cultured in 90% α-MEM (Gibco) with 10% FBS and 1% penicillin-streptomycin. OCI-AML3 was cultured in 80% α-MEM with 20% FBS and 1% penicillin-streptomycin. Lenti-X 293T cells were cultured in 90% DMEM with 10% FBS, 1% penicillin-streptomycin and 1% glutamine (Sigma-Aldrich). Cell line identities were verified by Short Tandem Repeats (STR) analysis. Primary MLL-AF6 patient samples were obtained via bone marrow aspiration at the Christie NHS Foundation Trust, following informed written consent in accordance with institutional guidelines and ethical approval. Primary patient cells were cultured in StemSpan SFEM medium (STEMCELL Technologies) with 1% penicillin-streptomycin, hFLT3L (100 ng/mL, PeproTech), hIL-3 (100 ng/mL, PeproTech), hTPO (100 ng/mL, PeproTech), hIL6 (100 ng/mL, PeproTech), hSCF (100 ng/mL, PeproTech), and UM729 (1 µM, STEMCELL Technologies). All cells were maintained at 37 °C in a humidified atmosphere with 5% CO₂ and were routinely tested and confirmed to be free of *Mycoplasma* contamination.

### Design and cloning of sgRNAs

Dual-sgRNA CRISPR interference libraries were cloned following a modified protocol from the Weissman lab^42^. The Perturb-seq vector pJR103 (Addgene #187242) was digested with BstXI (NEB) and BlpI (NEB) to create an insertion site for dual-sgRNA sequences. sgRNAs were designed using the Broad Institute’s GPP sgRNA Designer (https://portals.broadinstitute.org/gppx/crispick/public), and the sequences are listed in Supplementary Table 1. Complementary dual-sgRNA oligonucleotides containing BsmBI restriction sites were annealed, phosphorylated with T4 PNK (NEB), diluted, and ligated into the digested pJR103 backbone. An additional U6 promoter sequence was retrieved from pJR98 (Addgene #187239) using BsmBI-V2 (NEB), and cloned into the BsmBI-V2 site in the double sgRNA-containing pJR103 vectors. The plasmid pHR-UCOE-EF1a-ZIM3-dCas9-P2A-GFP (Addgene #188778) was used for the expression of dCas9-ZIM3-KRAB-GFP in CRISPR interference experiments.

LentiCRISPR v2-GFP (Addgene #82416) and LentiCRISPR v2-mCherry (Addgene #99154) vectors were used for CRISPR–Cas9-based gene knockout. Both vectors were digested with BsmBI-v2 (NEB) to create an insertion site for the sgRNA sequence. sgRNAs were designed using the Broad Institute’s GPP sgRNA Designer for CRISPR knockout (https://portals.broadinstitute.org/gppx/crispick/public). sgRNA sequences were inserted into the digested LentiCRISPR v2-GFP and v2-mCherry vectors by annealing complementary oligonucleotides, phosphorylation with T4 PNK (NEB), diluted, and ligation using T4 DNA ligase (NEB).

### Lentivirus generation and transduction

Lentiviral particles were produced in Lenti-X 293T cells using a second-generation packaging system. The transfer plasmid, pCMVΔ8.91, and pMD2.G were mixed at a 0.5:0.34:0.16 ratio with polyethyleneimine (PEI) and incubated at room temperature for 20 minutes before being added to the cells. Viral supernatant was collected at 48 and 72 hours post-transfection and filtered through a 0.45 µm PVDF filter (Starlab).

Target cells were seeded at appropriate densities 24 hours prior to infection. For transduction, cells were spinfected with virus mixed with 8 μg/mL polybrene (Bio-Techne) for 1.5 hours at 1,500 rpm at 32 °C. Following centrifugation, cells were incubated for 24 hours before the medium was replaced with fresh growth media.

### Flow cytometry

For cell sorting, human AML cells were washed with PBS containing 2% FBS and filtered through a 40 µm cell strainer to obtain single-cell suspensions. Mouse bone marrow cells were isolated by crushing femurs, tibias and pelvic bones with a mortar and pestle, following by filtration through a 40-μm cell strainer. Prior to sorting, mouse cells were stained on ice for 30mins with APC anti-CD45 (BD Biosciences). Target cell populations from both human and mouse samples were sorted using a BD FACS Aria III (BD Biosciences), and collected into tubes containing pre-chilled culture medium supplemented with 50% FBS and used for downstream applications.

For flow cytometry analysis, cells were incubated with combinations of the antibodies listed in Supplementary Table 3. Markers used to assess myeloid differentiation included PE/Cyanine7-conjugated anti-CD86 (BioLegend), and Brilliant Violet 711-conjugated anti-CD14 (BioLegend). Cells were stained with the antibody cocktail on ice for 30 minutes prior to acquisition. Apoptosis was measured using the eBioscience Annexin V Apoptosis Detection Kit (Invitrogen), following the manufacturer’s instructions, with Hoechst 33342 (Invitrogen) substituted for propidium iodide. Flow cytometry analyses were performed using a BD LSRFortessa™ (BD Biosciences) and all data analysed with FlowJo software.

### Perturb-seq

Primary patient cells were lentivirally transduced with ZIM3-KRAB-dCas9-GFP and pooled sgRNA library. 6 days of post-transduction, double positive (BFP^+^GFP^+^) cells were isolated by FACS. Sorted cells were washed and resuspended in PBS containing 0.04% BSA for single-cell encapsulation. Cell suspensions were loaded onto a Chromium Chip K (10x Genomics), and cDNA was prepared using the Chromium Next GEM Single Cell 5’ Kit v2 (10x Genomics) and the Chromium 5’ Feature Barcode Kit (10x Genomics), following the manufacturer’s instructions with the following modifications. As previously described^42^, a sgRNA-specific RT primer was added to the reverse transcription master mix. After cDNA amplification, size fractionation was performed according to the 10x Genomics protocol to separate the amplified cDNA (gene expression fraction) from the Perturb-seq sgRNA library. A subsequent 0.6× AMPure XP bead-based size selection was performed on the gene expression fraction to remove excess smaller material. Libraries were quality-checked using the Agilent Fragment Analyzer and quantified by qPCR using the KAPA Library Quantification Kit for Illumina Platforms (Roche). A low-output sequencing run was performed on the Verogen MiSeq FGx for quality control and sample balancing, followed by paired-end sequencing on an Illumina NovaSeq 6000.

### Cell proliferation assay

Lentivirally transduced cells were sorted by FACS and assessed for proliferation using the CellTiter-Glo Luminescent Cell Viability Assay (Promega), according to the manufacturer’s instructions. Cells were seeded in 96-well white-walled plates at a density of 500–1,000 cells per well, depending on the cell type. At the indicated time points, an equal volume of CellTiter-Glo reagent was mixed with the cell culture, and luminescence was measured using a microplate reader.

### Overexpression experiments

For the overexpression rescue experiments, an IRF2BP2 ORF with silent mutations in the protospacer adjacent motif (PAM) regions targeted by the sgRNAs was synthesized by GeneArt (Thermo Fisher Scientific) and cloned into the pKAM-BFP-Puro vector (Addgene #101864), which had been digested with BamHI and MluI (NEB). Cells were co-transduced with lentiviral constructs expressing either the IRF2BP2-targeting sgRNAs or non-targeting control sgRNAs, along with either a PAM-mutated IRF2BP2 ORF or an empty vector. Following transduction, cellular differentiation and apoptosis were assessed by flow cytometry.

### Animal experiments

NSGS (NOD.Cg-*Prkdc*^scid^ *Il2rg*^tm1Wjl^ Tg(CMV-IL3,CSF2,KITLG)1Eav/MloySzJ) mice were purchased from The Jackson Laboratory. All mouse work was performed in accordance with the United Kingdom Animals (Scientific Procedures) Act 1986, and were reviewed and approved by the Animal Welfare and Ethical Review Body (AWERB) at the Cancer Research UK Manchester Institute (CRUK-MI). GFP⁺mCherry⁺ double-positive cells were sorted using a BD FACSAria III. A total of 5 × 10⁴ sorted cells per mouse were resuspended in sterile PBS and injected via tail vein into sublethally irradiated (100 cGy) NSGS mice. Irradiation was performed 4–6 hours prior to transplantation. Mice were monitored regularly for signs of disease or distress. Bone marrow was harvested for analysis when mice exhibited signs of ill health.

### Western blot analysis

Cells were lysed in RIPA buffer (Sigma-Aldrich) supplemented with 1% protease inhibitor cocktail (Sigma-Aldrich) for 20 minutes at 4 °C, followed by centrifugation at 14,000 x g to collect the protein-containing supernatant. Proteins were prepared for separation using NuPAGE Sample Reducing Agent (Invitrogen) and NuPAGE LDS Sample Buffer (Invitrogen), incubated at 70 °C for 10 minutes, and separated on NuPAGE 4%–12% Bis-Tris polyacrylamide gels (Invitrogen), followed by transfer to nitrocellulose membranes using iBlot® Gel Transfer Stacks (Invitrogen). Membranes were blocked for 1 hour at room temperature, then incubated overnight at 4 °C with primary antibodies according to the manufacturers’ instructions. The following antibodies were used: anti-IRF2BP2 (Proteintech), anti-β-actin (Sigma-Aldrich), anti-DNMT1 (Cell Signaling), and anti-TRIM28 (Invitrogen). Proteins were visualized using either Rabbit IgG (H+L) (invitrogen) or mouse IgG (H+L) (invitrogen), with ECL prime western blotting detection reagent (Cytiva). Imaging was performed on a ChemiDoc Imaging system (Bio-Rad).

### Rapid immunoprecipitation mass spectrometry of endogenous proteins (RIME)

20 μg of anti-IRF2BP2 antibody (Proteintech) or IgG control antibody (Abcam) were incubated with Protein A magnetic Dynabeads (Invitrogen) per reaction, overnight. 1 × 10⁸ THP-1 cells per sample were cross-linked with 1% formaldehyde (Sigma-Aldrich) for 10 minutes, followed by quenching with 0.125 M glycine (Sigma-Aldrich) for 5 minutes. Cells were then incubated in lysis buffer 1 (50 mM HEPES-KOH, pH 7.5; 140 mM NaCl; 1 mM EDTA; 10% glycerol; 0.5% NP-40; 0.25% Triton X-100; and 1% protease inhibitors) for 10 minutes at 4 °C, followed by incubation in lysis buffer 2 (10 mM Tris-HCl, pH 8.0; 200 mM NaCl; 1 mM EDTA; 0.5 mM EGTA; and 1% protease inhibitors) for another 10 minutes at 4 °C. After centrifugation at 2,000 × g for 5 minutes at 4 °C, the pellet was resuspended in lysis buffer 3 (10 mM Tris-HCl, pH 8.0; 200 mM NaCl; 1 mM EDTA; 0.5 mM EGTA; 0.1% sodium deoxycholate; 0.5% N-lauroylsarcosine; and 1% protease inhibitors) and subjected to sonication using a Diagenode Bioruptor at 4 °C (8 cycles of 30 seconds on and 30 seconds off).

Sonicated nuclei were enriched by the addition of 10% Triton X-100, followed by centrifugation at 14,000 x g speed for 10 minutes at 4 °C. Fragment size was verified by agarose gel electrophoresis, ensuring DNA fragments between 200–600 bp. The resulting supernatants were incubated overnight at 4 °C with antibody-conjugated magnetic beads. The following day, the beads were washed eight times with RIPA buffer (50 mM HEPES-NaOH, pH 7.6; 500 mM LiCl; 1 mM EDTA; 1% NP-40; 0.7% sodium deoxycholate; and 1% protease inhibitors). Proteins were digested with trypsin (Sigma-Aldrich) at 37 °C for 18 hours. Peptides were analysed using an Ultimate 3000 RSLCnano system (Thermo Scientific) coupled to an Orbitrap Lumos mass spectrometer (Thermo Scientific). Data analysis was performed using Mascot Distiller and Mascot (Matrix Science) against the human protein database.

### Duolink Proximity Ligation Assay

Duolink Proximity Ligation Assay was performed using the Duolink In Situ Red Starter Kit Mouse/Rabbit (Sigma-Aldrich) according to the manufacturer’s instructions. 5 × 10⁴ THP-1 cells per reaction were cytospin onto glass slides, fixed in 4% paraformaldehyde for 15 minutes at room temperature, and permeabilized with 0.05% Triton X-100 in PBS for 5 minutes. Following washing, samples were blocked with Duolink blocking solution for 1 hour at 37 °C in a humidified chamber. Primary antibodies against IRF2BP2 (Proteintech), DNMT1 (Cell Signaling), and TRIM28 (Invitrogen) were diluted in Duolink Antibody Diluent, applied to the samples, and incubated overnight at 4 °C. The next day, after washing, Duolink PLA probes (PLUS and MINUS) were added and incubated for 1 hour at 37 °C. The ligation solution was then applied and incubated for 30 minutes at 37 °C, followed by additional washes.

The amplification solution was freshly prepared and added to the samples, which were incubated for 100 minutes at 37 °C in a pre-heated humidity chamber. After final washes, samples were mounted using Duolink® In Situ Mounting Medium with DAPI and imaged on a Zeiss LSM880 confocal microscope with Airyscan. PLA signals were quantified using FIJI (ImageJ) by analysing the number of dots per nucleus or per cell.

### RNA-seq

Total RNA was extracted from sorted cells 3 days post-transduction using the RNeasy Plus Mini Kit (Qiagen), and its concentration was measured using a NanoDrop spectrophotometer. cDNA libraries were prepared using the NEBNext Ultra II Directional RNA Library Prep Kit for Illumina (New England Biolabs) with the NEBNext Poly(A) mRNA Magnetic Isolation Module (New England Biolabs). Libraries were quantified by qPCR using a KAPA Library Quantification Kit for Illumina. Paired end 100bp sequencing was performed on a NovaSeq 6000 sequencer (Illumina).

### CUT&RUN

For each reaction, 400 × 10³ sorted cells (three days post-transduction) were pelleted and cross-linked with 1% formaldehyde in RPMI-1640 medium for 1 minute, then quenched with glycine to a final concentration of 125 mM. The cross-linked cells were washed twice with wash buffer (20 mM HEPES pH 7.9, 150 mM NaCl, 1% Triton X-100, 0.05% SDS, 1× PIC, 0.5 mM spermidine), incubated with 11 μL of Concanavalin A beads (Bangs Laboratories) for 10 minutes at room temperature, followed by incubation with 0.5 μL of antibody overnight (TRIM28 [Abcam], IRF2BP2 [Sigma-Aldrich], H3K9me3 [Abcam], IgG [Abcam]). Each reaction was washed twice with wash buffer containing 0.01% digitonin, followed by incubation (1:300) with pAG-MNase (a gift from the Hurd laboratory, QMUL) for 10 minutes at room temperature. After two washes, digestion was activated by adding 50 μL of digitonin-wash buffer supplemented with 2 mM CaCl₂ and incubating for 2 hours at 4 °C. Digestion was stopped with 33 μL of stop buffer (340 mM NaCl, 20 mM EDTA, 4 mM EGTA, 50 μg/mL RNase A, 50 μg/mL glycogen). Following a 10-minute incubation at 37 °C, the Concanavalin A beads and supernatant were separated, and 0.8 μL of 10% SDS and 1 μL of 20 μg/μL Proteinase K (Invitrogen) were added to the supernatant and incubated overnight at 55 °C. DNA was extracted using the DNA Clean & Concentrator-5 Kit (Zymo Research), and libraries were prepared using the NEBNext® Ultra™ II DNA Library Prep Kit for Illumina (New England Biolabs). Final library quality was assessed using the Agilent 4200 TapeStation System with D1000 ScreenTapes. Sequencing was performed on an Illumina NovaSeq X Plus.

### Transfection

Cells were transfected using the Neon Transfection System (Thermo Fisher Scientific) according to the manufacturer’s instructions. Cells were seeded at an appropriate density one day prior to transfection to achieve 70–80% confluency. After washing with PBS, cells were resuspended in Neon Resuspension Buffer R. A total of 5 × 10⁶ cells were mixed with 5 µg of PiggyBac transposon plasmid and PiggyBac transposase expression plasmid at a 1:1 ratio. Electroporation was performed using the following settings: 1200 V, 20 ms, 2 pulses. After electroporation, cells were transferred to pre-warmed complete growth medium and incubated for 2 days prior to FACS to isolate transfected cells.

### CRISPR activation and CRISPR interference

For the CRISPR activation experiments, cells were transfected with vector encoding either LTR5_Hs S. aureus scaffold CARGO array (Addgene # 191317) or LTR5_Hs S. pyogenes scaffold CARGO array (Addgene # 191316). Transfected mcherry^+^ cells were sorted using FACS and expanded for a week. Cells were then lentivirally transduced with dCAS9-VP64-GFP (Addgene # 61422), following by functional assays.

To assess whether repression of LTR5_Hs could rescue the effects of IRF2BP2 knockdown, cells were first transfected with either the LTR5_Hs *Staphylococcus aureus* scaffold CARGO array (Addgene #191317) or the LTR5_Hs *Streptococcus pyogenes* scaffold CARGO array (Addgene #191316). mCherry⁺ cells were enriched by FACS and expanded for one week. The sorted cells were then transduced with a GFP-tagged dCas9-ZIM3-KRAB construct and BFP-labelled sgRNAs targeting either *IRF2BP2* or a NT control, and subsequently used for downstream phenotypic assays.

### RT–qPCR

Total RNA from sorted target cells was isolated using the RNeasy Plus Mini Kit (Qiagen). Complementary DNA (cDNA) was synthesized using the High-Capacity cDNA Reverse Transcription Kit (Applied Biosystems), according to the manufacturer’s instructions. Quantitative PCR (qPCR) was performed using the FastStart Universal SYBR Green Master (Roche) on a QuantStudio 3 Real-Time PCR System (Applied Biosystems).

### Perturb-seq analysis

Transcript quantification and alignment was performed using CellRanger (v9.0.0) with reference genome GRCh38. CRISPR guides were quantified using CellRanger Multi. Subsequent analyses were implemented using Scanpy (v1.10.3). scRNA-seq cells were filtered by removing doublets (Scrublet, 0.2.3), and low-quality cells were filtered according to recommended best practices guidelines (https://doi.org/10.1038/s41576-023-00586-w). Cells with fewer than 1000 counts or 500 genes detected were also removed. CRISPR guide counts were processed by removing all cells with fewer than 5 guide counts and then assigning the most abundant guide identity to each cell. The cells with high counts of multiple guides were also removed as possible doublets. Two NT control groups were combined for downstream analysis. To confirm the efficacy of each CRISPR guide, we performed a T-test (Scanpy, v1.10.3) comparing the target gene and non-targeting control cells, using 0.05 as the significance threshold.

## Mixscape Analysis

To identify cells with successful perturbations, we used the Pertpy (v0.9.4) implementation of Mixscape. Local perturbation signatures and subsequent dimensionality reduction were calculated as per the default parameters. The Mixscape model was then used (logfc_threshold = 0.15, min_de_genes=5) to assign Mixscape class identities. LDA was used to visualise the KO cells relative to the NT control. Differential gene expression (Scanpy, v1.10.3) was then used to compare the KO cells to NT control cells with a significance threshold of 0.05.

### RNA-seq analysis

Raw sequencing reads underwent adapter trimming and quality filtering using fastp (v0.23.4). Cleaned reads were aligned to the human reference genome (hg38) using STAR (v2.7.10a). BAM files were merged, sorted, and indexed using samtools (v1.17). For standard gene expression analysis, only uniquely mapped reads were retained, and transcript abundance was quantified using featureCounts (v2.0.6). For transposable element analysis, multimapping reads were retained, and quantified using the TEtranscripts (v2.2.3). Differential expression analysis for both gene and TE datasets was performed using the DESeq2 package in R. Gene Ontology (GO) enrichment and Gene Set Enrichment Analysis (GSEA) were conducted using the clusterProfiler package in R.

### CUT&RUN analysis

CUT&RUN sequencing reads were quality-checked using FastQC (v0.12.1) and trimmed for adapter sequences and low-quality bases using Trim Galore (v0.6.10). Cleaned reads were aligned to the human reference genome (hg38) using Bowtie2 (v2.5.1), and the resulting BAM files were sorted and indexed using SAMtools (v1.17). PCR duplicates were removed using Picard MarkDuplicates (v3.0.0).

Multimapping reads were retained to enable downstream analysis of transposable elements. Genome coverage files were generated using BEDTools genomecov (v2.31.0) and converted to BigWig format using bedGraphToBigWig (v20230927, UCSC tools). Peak regions were visualized using deepTools (v3.5.2) for heatmap generation. Transposable elements were annotated based on RepeatMasker annotations.

### Statistical analysis

Statistical analyses were performed using python, R (4.4.1) and GraphPad Prism 10. Comparisons were conducted using unpaired two-tailed t-test, one-way ANOVA or two-way ANOVA. P values of less than 0.05 were considered significant. Errors bars represented standard deviation. Graphs and visualizations were generated using python, GraphPad Prism and R packages.

### Data availability

All data generated to support the findings of this study are available. The raw and processed RNA-seq, CUT&RUN, and Perturb-seq datasets have been deposited in the GEO. Publicly available RNA-seq data were obtained from the GEO under accession number GSE135207.

## Acknowledgements

We thank Wolfgang Breitwieser, John Weightman, Rachel Horner, Hannah Bowden, and Jason Rumley from the CRUK-MI Molecular Biology Facility for assistance with next-generation sequencing; Yvonne Connolly, Robert Faulkner, and Adam Flinders from the Biological Mass Spectrometry Facility for assistance with mass spectrometry; Antonia Banyard, Yosra Elagili, Emily Scanlon, and Adam Milner from the Flow Cytometry Facility for assistance with flow cytometry; Sudhakar Sahoo, Anoop Sanalkumar, Stephen Kitcatt from the Computational Biology and Scientific Computing Facilities and Yitao Chen from the National Biomarker Centre for assistance with data analysis; Natalia Moncaut and Lauren Street from the Genome Editing and Mouse Models Facility for assistance with the mouse model; and Steve Bagley and Henry Banks from the Visualisation, Irradiation and Analysis Facility for assistance with imaging. We are also grateful to all members of the CRUK MI Biological Resources Unit for their care and processing of mice used in this study. This work was supported by Cancer Research UK (C5759/A20971 (G.L.) and C5759/A27412 (G.L.).

## Author information

J.X. conceived the study, designed and performed experiments, analyzed data, and wrote the manuscript. J.W. performed data analysis. L.C. performed experimental work. M.L. contributed to manuscript writing and experimental design. E.A.L., performed experimental work and contributed to data analysis. R.S. provided support with data analysis. O.D. supervised E.A.L.’s work and provided guidance on related experiments and manuscript preparation. G.L. supervised the overall study, provided funding and wrote the manuscript.

---

**Extended Data Fig. 1.**
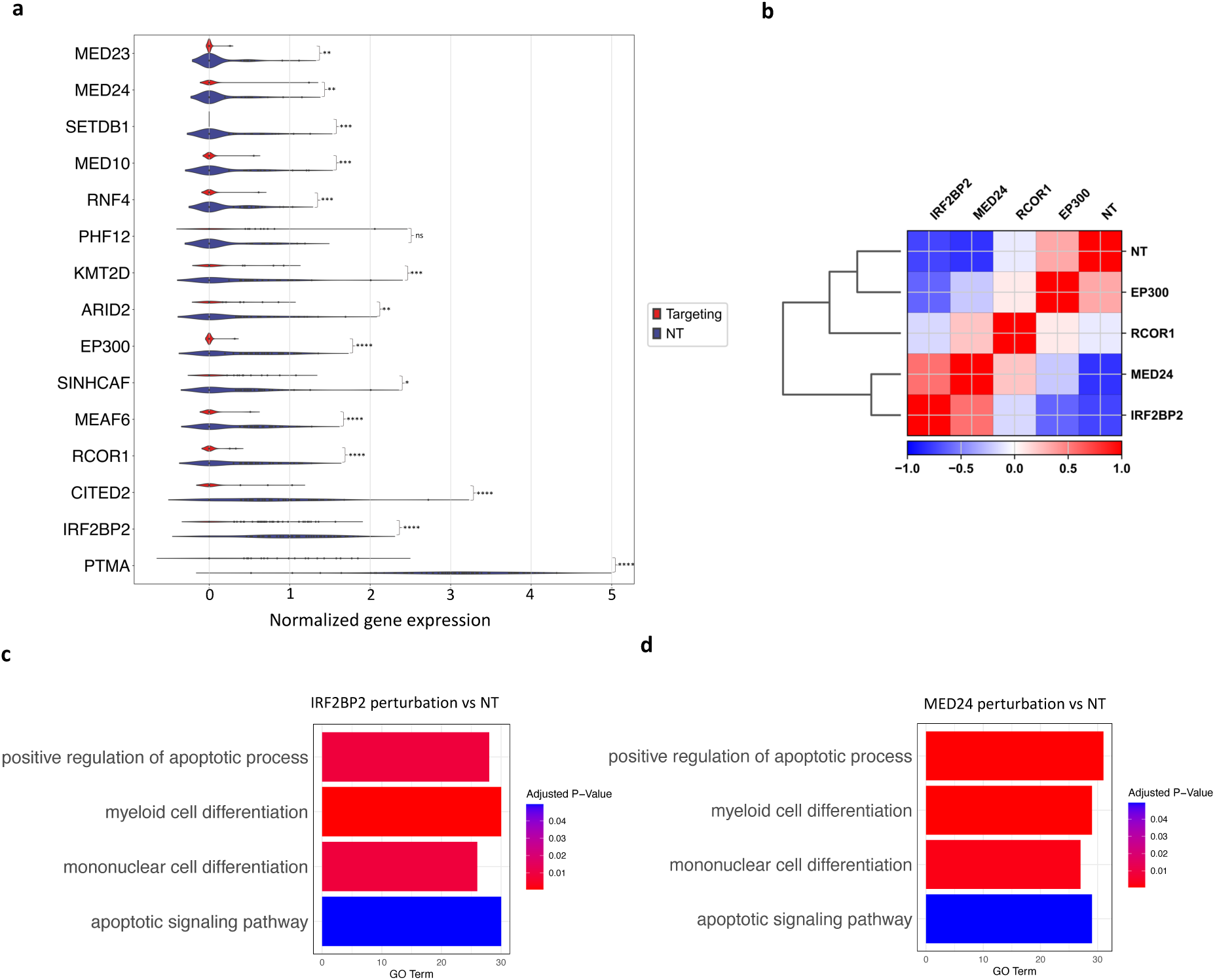
Perturb-seq reveals the key chromatin-associated regulators in AML. **a**, Violin plot showing normalised gene expression levels (x-axis) of the 15 targeted chromatin-associated regulators (y-axis) analyzed by single-cell perturb-seq in the presence of either the control NT sgRNA (blue) or the targeting sgRNAs (red). All targeted genes within each perturbation group were significantly downregulated, except for *PHF12*. Statistical test: two-sided unpaired t-test. **b**, Correlation coefficients between the mean profiles of cells under each perturbation condition. Red indicates a stronger correlation coefficient, while blue indicates a weaker correlation coefficient. **c,d**, Bar plot showing Gene Ontology (GO) enrichment analysis of IRF2BP2 (**c**) and MED24 (**d**) KD. * *P* < 0.05; ** *P* < 0.01; *** *P* < 0.001; **** *P* < 0.0001.

**Extended Data Fig. 2.**
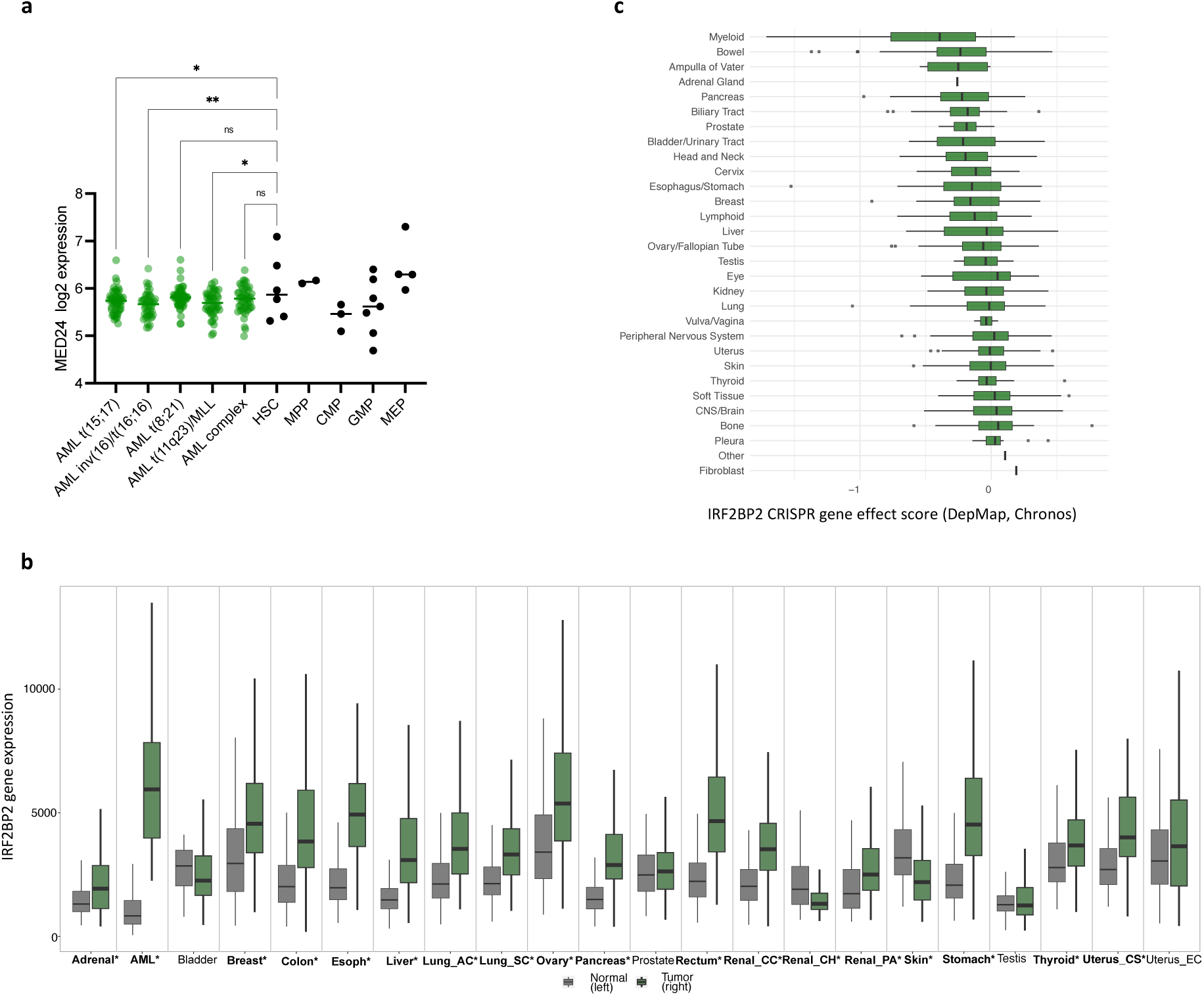
*MED24* expression in hematopoietic cells and *IRF2BP2* expression across cancers and solid tissues. **a**, Gene expression levels of *MED24* were retrieved from the BloodSpot database. One-way ANOVA. **b**, Bar plot comparing *IRF2BP2* expression between cancers (green bar, right) and corresponding tissues (grey bar, left), obtained from the TNMplot database. An asterisk (*) indicates a significant difference between cancer and corresponding normal tissues. Mann–Whitney U test. **c**, Bar plot showing the average CRISPR gene effect score of IRF2BP2 across various cancer types (DepMap). More negative scores indicate greater dependency. * *P* < 0.05; ** *P* < 0.01.

**Extended Data Fig. 3.**
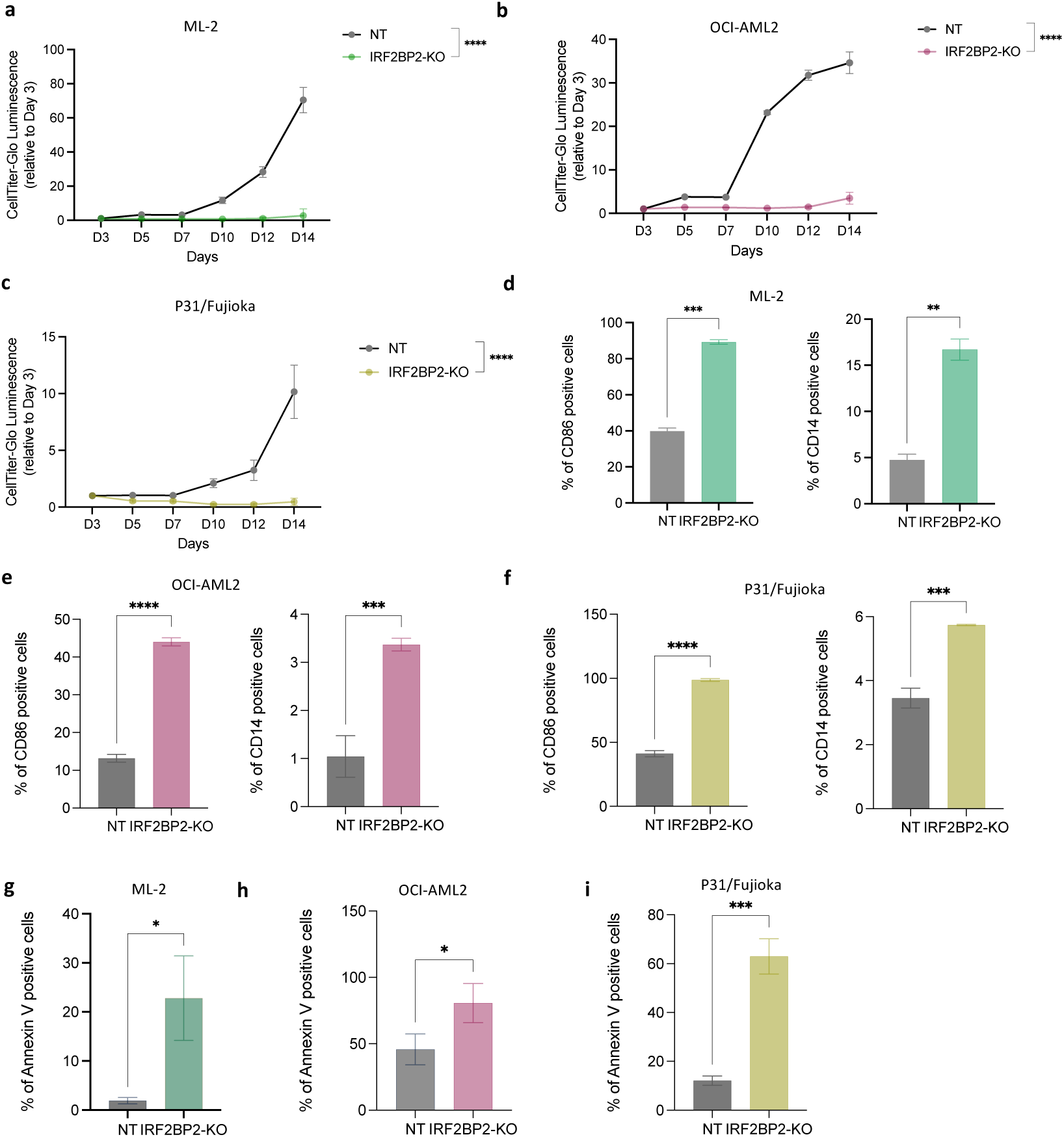
IRF2BP2 is a shared dependency among multiple AML subtypes. a–c. Cell proliferation time course, as measured by CellTiter-Glo, for ML-2 (**a**), OCI-AML2 (**b**) and P31/Fujioka (**c**) cells following CRISPR/Cas9-mediated KO of IRF2BP2. Transduced cells were sorted based on GFP and mCherry reporters at day 3. Statistical test: Two-way ANOVA. **d-f**, Expression of the differentiation markers CD86 and CD14 was assessed by flow cytometry in ML-2 (**d**), OCI-AML2(**e**), and P31/Fujioka (**f**) cells after 7 days transduction. Unpaired t-test. **g-i**, ML-2 (**g**), OCI-AML2 (**h**), and P31/Fujioka (**i**) cells were analyzed by flow cytometry 10 days post-transduction following Annexin V and Hoechst staining to evaluate apoptotic cell populations. Unpaired t-test. Values are expressed as mean ± s.e.m (n = 3). * *P* < 0.05; ** *P* < 0.01; *** *P* < 0.001; **** *P* < 0.0001.

**Extended Data Fig. 4.**
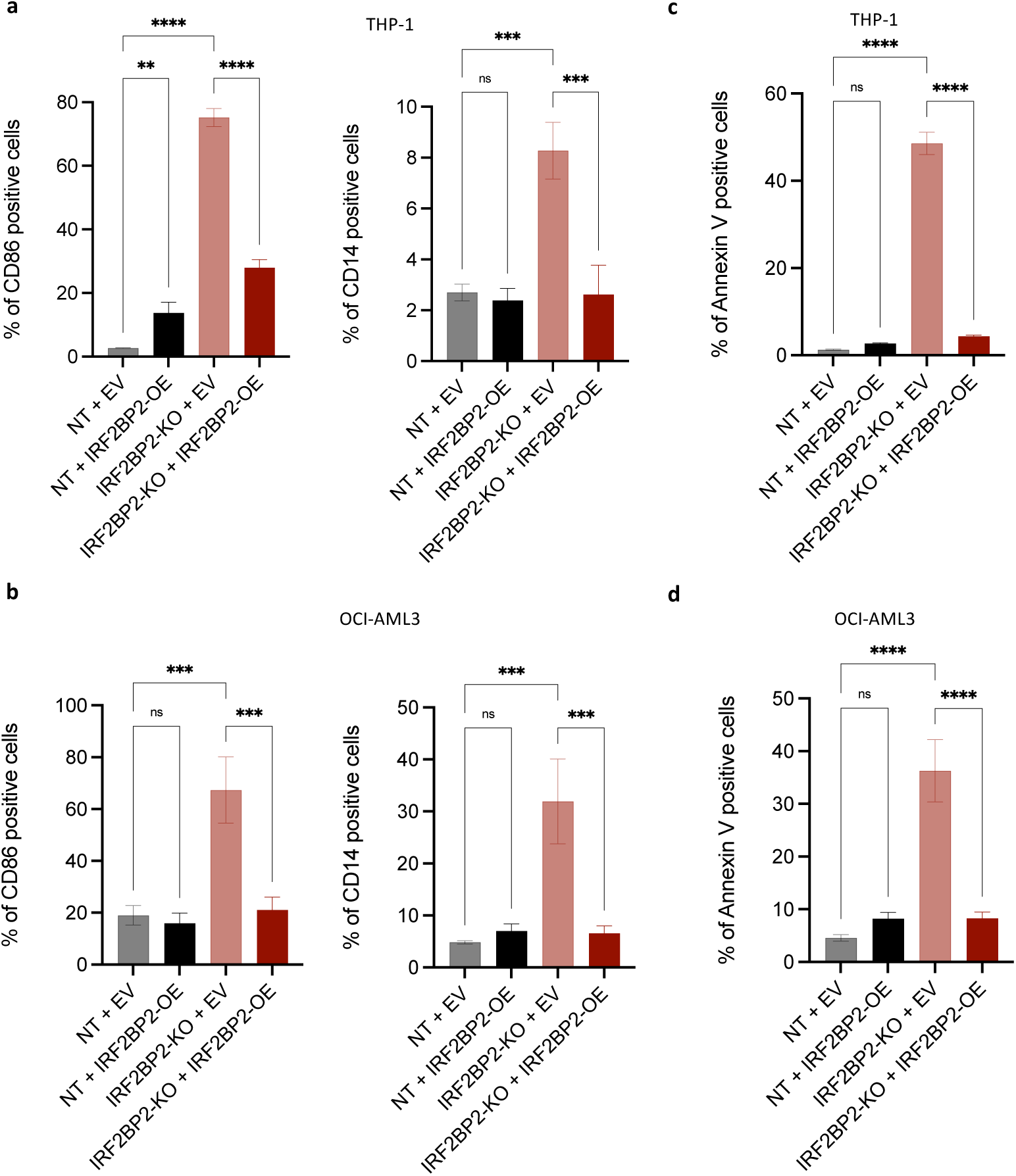
Ectopic expression of IRF2BP2 reverses the phenotypic consequences of endogenous IRF2BP2 KO. a,b, THP-1 and OCI-AML3 cells were transduced to overexpress the IRF2BP2 open reading frame (IRF2BP2-OE) or an empty vector (EV), along with either IRF2BP2-targeting sgRNAs or NT sgRNAs. Flow cytometry for myeloid differentiation markers was performed in THP-1 (**a**) and OCI-AML3 (**b**) cells at 7 days post-transduction. **c,d**, Apoptosis assay in THP-1 (**c**) and OCI-AML3 (**d**) cells was performed at 10 days post-transduction. Data are shown as mean ± s.e.m (n = 3), one-way ANOVA. ** *P* < 0.01; *** *P* < 0.001; **** *P* < 0.0001.

**Extended Data Fig. 5.**
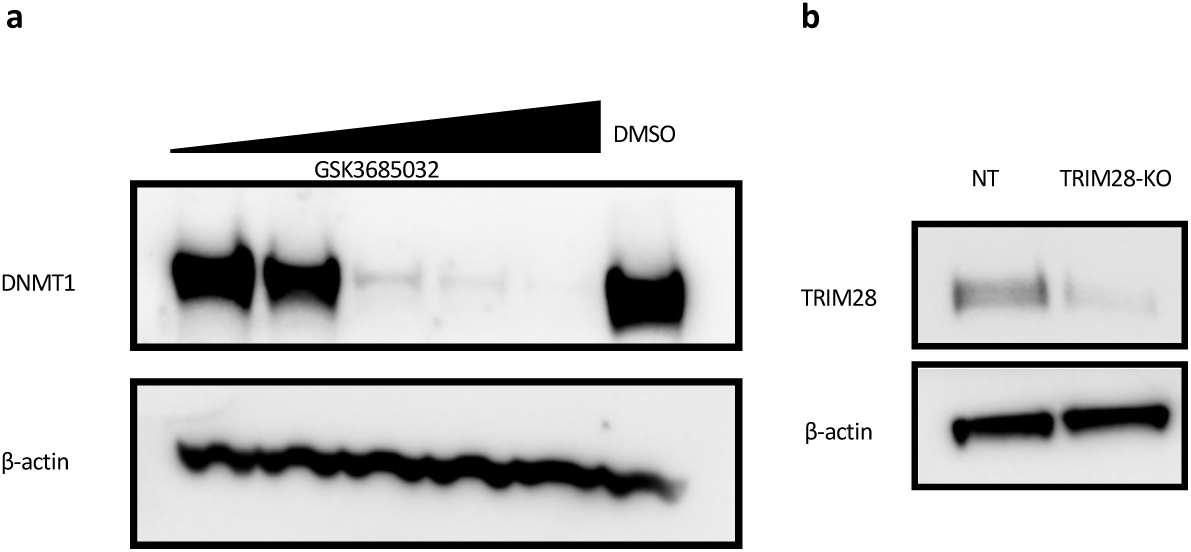
Validation of DNMT1 and TRIM28 depletion. **a**, Western blot analysis of DNMT1 protein levels in THP-1 cells treated for 2 days with increasing concentrations of the DNMT1 inhibitor GSK3685032 (0nM, 4nM, 40nM, 400nM, and 4μM) or with DMSO. b, Western blot analysis validating TRIM28 loss in TRIM28-KO THP-1 cells. β-actin was used as a loading control.

**Extended Data Fig. 6.**
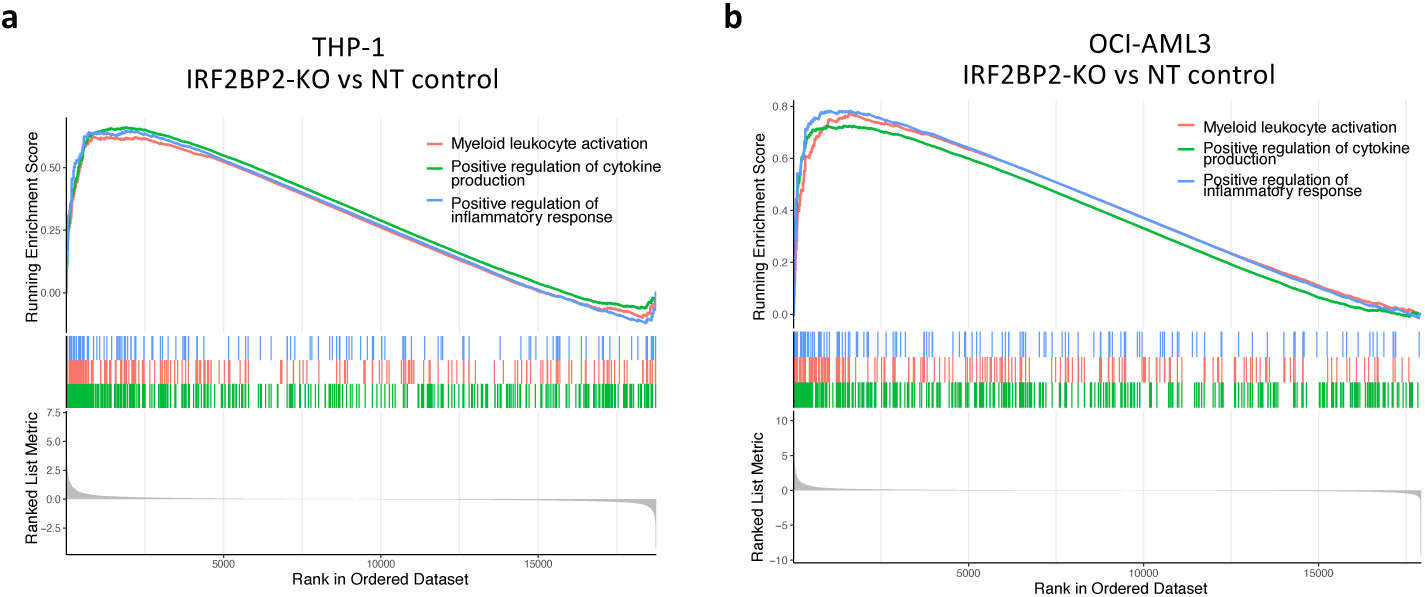
IRF2BP2 is required for suppressing immune response signaling. a,b, Gene Set Enrichment Analysis (GSEA) plots showing enrichment of selected upregulated gene sets in IRF2BP2-KO THP-1 (**a**) and OCI-AML3 (**b**) cells compared to NT controls.

**Extended Data Fig. 7.**
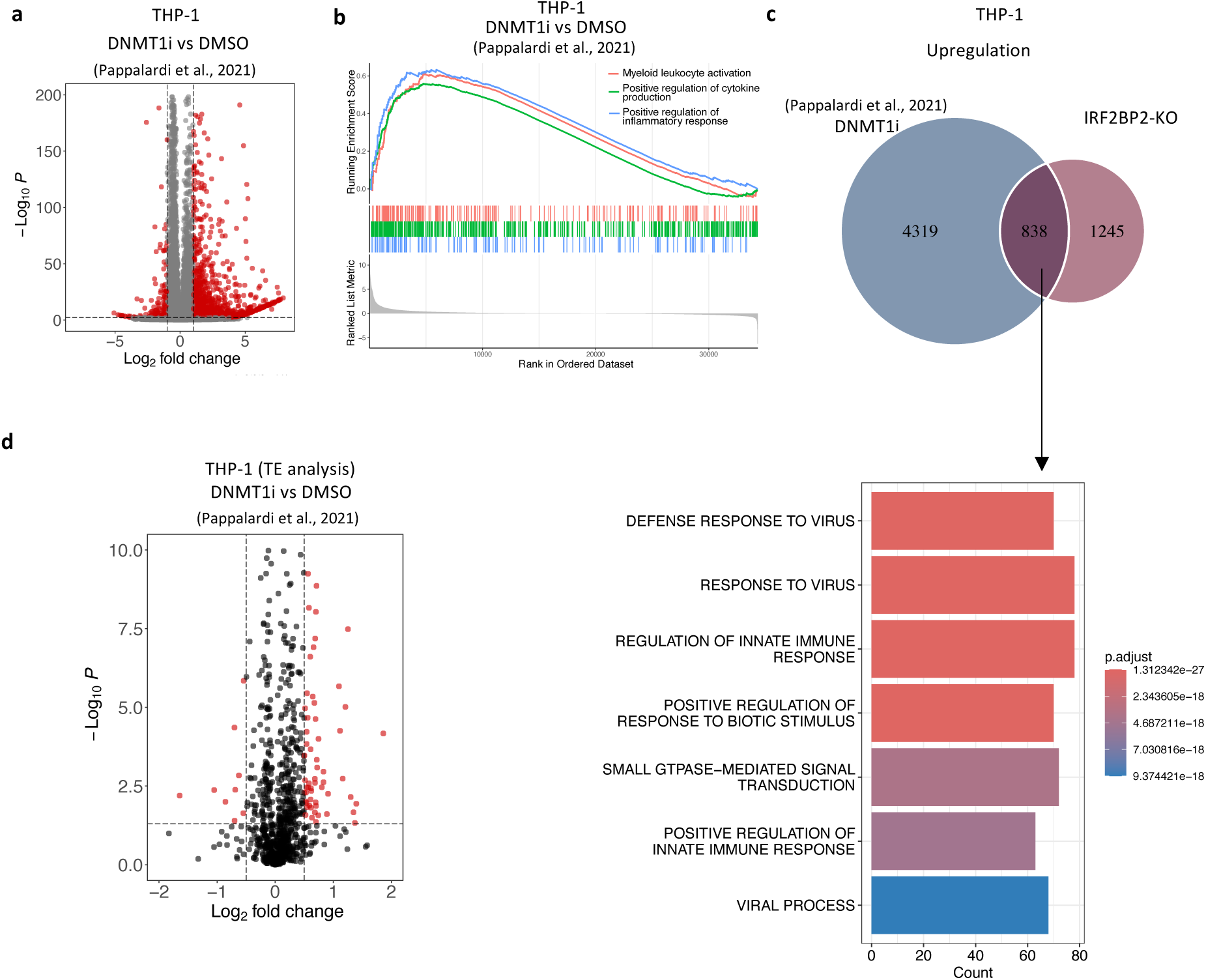
Loss of DNMT1 recapitulates the transcriptional changes observed upon IRF2BP2 depletion. a, RNA-seq analysis of THP-1 cells following GSK3685032 treatment (400 nM, 4 days) using publicly available data^27^. Volcano plot displays differentially expressed genes compared to vehicle-treated controls. Significantly upregulated and downregulated genes (P < 0.05, |log_₂_ fold change| > 1) are highlighted in red. **b**, GSEA plot illustrating the enrichment of selected upregulated gene sets in THP-1 cells treated with GSK3685032 relative to controls. **c**, Venn diagram showing the overlap of significantly upregulated genes (adjusted P < 0.05) between IRF2BP2-KO cells (three days post-transduction) and THP-1 cells treated with DNMT1 inhibitor GSK3685032 (400 nM, 4 days). GO enrichment analysis of genes commonly upregulated in both IRF2BP2-KO cells and GSK3685032-treated THP-1 cells. **d**, RNA-seq analysis of transposable elements in THP-1 cells treated with GSK3685032 (400 nM, 4 days). Volcano plot depicting differential transposable expression in THP-1 cells following GSK3685032 treatment. Significantly upregulated and downregulated genes (P < 0.05, |log_₂_ fold change| > 0.5) are highlighted in red.

**Extended Data Fig. 8.**
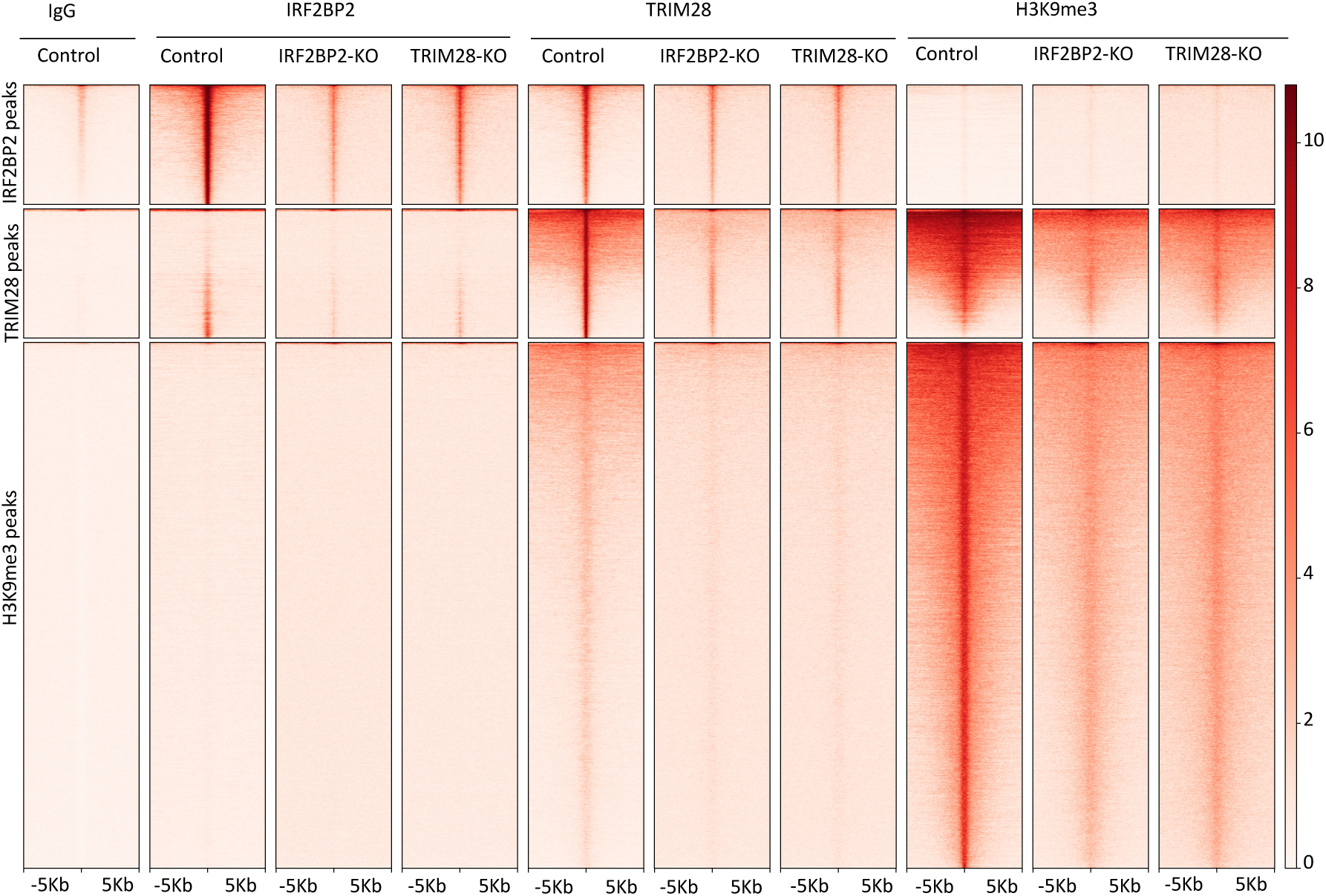
IRF2BP2 loss reduces TRIM28 and H3K9me3 occupancy at non-repetitive genomic regions. Heatmaps of CUT&RUN signal for IRF2BP2, TRIM28, and H3K9me3 across the IRF2BP2-, TRIM28-, and H3K9me3-defined peak sets in control, IRF2BP2-KO, and TRIM28-KO THP-1 cells.

**Extended Data Fig. 9.**
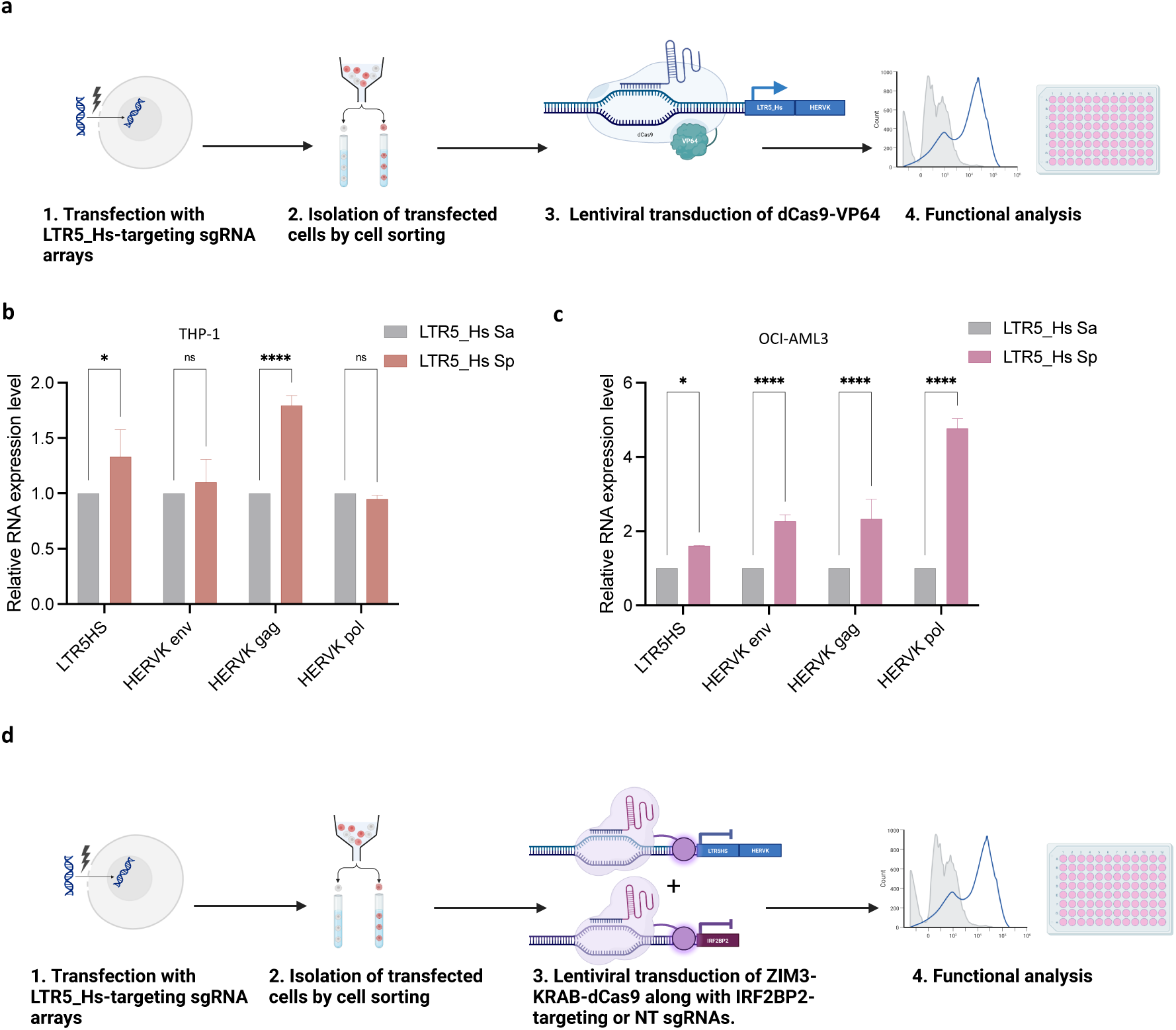
CRISPR-mediated activation/repression of HERVK/LTR5_Hs in AML. **a**, Schematic diagram illustrating the workflow of CRISPR activation of HERVK/LTR5_Hs in AML. **b,c,** RT-qPCR analysis showing increased expression of LTR5_Hs and HERVK elements (env, gag and pol) following CRISPR activation in THP-1 (**b**) and OCI-AML3 (**c**) cells. Two-way ANOVA. Data shown as mean ± s.e.m (n = 3). * *P* < 0.05; **** *P* < 0.0001. **d,** Schematic of the rescue experiment using CRISPR-mediated interference to simultaneously target HERVK/LTR5_Hs and IRF2BP2.

**Extended Data Fig. 10.**
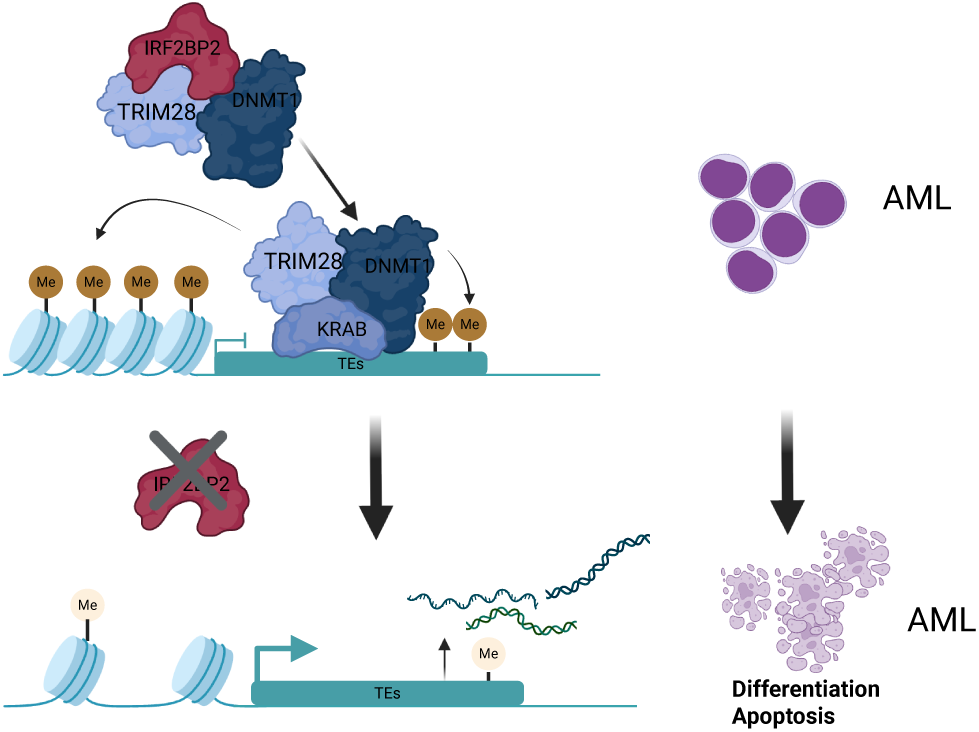
IRF2BP2 is a selective dependency of AML through repression of transposable elements. Schematic of the working model: IRF2BP2 maintains AML survival by epigenetically repressing TE expression through its interaction with TRIM28 and DNMT1.

